# Shedding dynamics of a DNA virus population during acute and long-term persistent infection

**DOI:** 10.1101/2025.03.31.646279

**Authors:** Sylvain Blois, Benjamin M. Goetz, Anik Mojumder, Christopher S. Sullivan

**Affiliations:** Department of Molecular Biosciences, LaMontagne Center for Infectious Disease, The University of Texas at Austin, Austin, Texas, 78712, USA; Department of Biomedical Sciences, University of Cagliari, Monserrato, Cagliari, 09042, Italy; Center for Biomedical Research Support, The University of Texas at Austin, Austin, Texas, 78712, USA

## Abstract

Although much is known of the molecular mechanisms of virus infection within cells, substantially less is understood about within-host infection. Such knowledge is key to understanding how viruses take up residence and transmit infectious virus, in some cases throughout the life of the host. Here, using murine polyomavirus (muPyV) as a tractable model, we monitor parallel infections of thousands of differentially barcoded viruses within a single host. In individual mice, we show that numerous viruses (>2600) establish infection and are maintained for long periods post-infection. Strikingly, a low level of many different barcodes is shed in urine at all times post-infection, with a minimum of at least 80 different barcodes present in every sample throughout months of infection. During the early acute phase, bulk shed virus genomes derive from numerous different barcodes. This is followed by long term persistent infection detectable in diverse organs. Consistent with limited productive exchange of virus genomes between organs, each displays a unique pattern of relative barcode abundance. During the persistent phase, constant low-level shedding of typically hundreds of barcodes is maintained but is overlapped with rare, punctuated shedding of high amounts of one or a few individual barcodes. In contrast to the early acute phase, these few infrequent highly shed barcodes comprise the majority of bulk shed genomes observed during late times of persistent infection, contributing to a stark decrease in bulk barcode diversity that is shed over time. These temporally shifting patterns, which are conserved across hosts, suggest that polyomaviruses balance continuous transmission potential with reservoir-driven high-level reactivation. This offers a mechanistic basis for polyomavirus ubiquity and long-term persistence, which are typical of many DNA viruses.

**Author Summary / Importance:** Polyomavirus infections, mostly benign but potentially fatal for immunocompromised individuals, undergo acute and long-term persistent infections. Typically, polyomavirus-associated diseases arise due to virus infection occurring in the context of a persistently infected individual. However, little is understood regarding the mechanisms of how polyomaviruses establish, maintain, and reactivate from persistent infection. We developed a non-invasive virus shedding assay combining barcoded murine polyomavirus, massively parallel sequencing technology, and novel computational approaches to track long-term infections in mice. We expect these methods to be of use not only to the study of DNA viruses but also for understanding persitent infection of diverse microbes. The study revealed organ-specific virus reservoirs and two distinct shedding patterns: constant low-level shedding of numerous barcodes and episodic high-level shedding of few barcodes. Over time, the diversity of shed barcodes decreased substantially. These findings suggest a persistent low-level infection in multiple reservoirs, with occasional bursts of replication in a small subset of infected cells. This combination of broad reservoirs and varied shedding mechanisms may contribute to polyomavirus success in transmission and maintaining long-term infections.

## Introduction

Fundamental questions about infection with most viruses remain unanswered, for example: 1) How many viruses establish an infection? 2) Is the virus diversity maintained as the infection progresses? 3) If a persistent infection ensues with ongoing shedding, are the shedders distributed evenly among the entire population or do they represent a subset? 4) What fraction of the virus population undergoes low-level continuous shedding (smoldering) and/or latent/lytic infections? 5) How do different tissues contribute to these processes? The advent of cost-effective high-throughput massively parallel sequencing and DNA barcoding technologies provides an opportunity to address these questions if suitable models can be employed.

Models of acute virus infection are prevalent due to their ease of study and relevance to disease. Genetically barcoded RNA virus populations have been engineered to study the effect of population bottlenecks, dissemination, mutations, and different routes of inoculation on virus population dynamics during acute infection (1–11). Less studied are the mechanisms of how viruses undergo and reactivate from persistent infections (12, 13). This is due, in part, to the challenges inherent in longitudinal infection studies *in vivo*.

Unlike most RNA viruses, many DNA viruses establish long-term persistent infection as an inherent component of their lifecycle (14, 15). To achieve this, it is widely accepted that either a cycle of latent/lytic or smoldering infections occur where viruses are never fully suppressed by the immune system (16). Latent/lytic infectious cycles result from reversible differential gene expression programs where high levels of viruses are sporadically produced from a limited number of cellular reservoirs only during the lytic phase (17). Smoldering infections occur with new viruses continuously replicating at low levels (16). For many DNA viruses, how and whether latent/lytic and/or smoldering infections occur remains poorly characterized. An ideal experimental system to address this requires tractable virus genetics in cultured cells and *in vivo*, along with the ability to produce genetically tagged viruses with little cost to viral fitness. Optimally, such a system would have the ability to undergo both acute and persistent infections and allow for non-invasive tracking of multiple virus lineages in a single experiment.

Polyomaviruses (PyVs) have been important laboratory models contributing to understanding fundamental principles of cell and molecular biology, the immune response, and cancer (18–24). PyVs are small DNA viruses with ∼5Kb circular genomes that undergo long-term persistent infections. Most humans are persistently infected with different species of PyVs (25) and previous studies identified kidneys as one of the persistent reservoirs of multiple PyVs in humans and mice (22). PyV infections emerging from persistence reservoirs are associated with several life-threatening diseases in immunocompromised patients (25–33). Although poorly understood, determining the mechanism of PyV persistence may lead to new approaches for preventing and treating PyV-associated diseases (25–33).

Previous studies showed that high levels of infectious murine PyV (muPyV) are excreted in the urine of healthy mice during the acute and persistent phases of infection (34, 35). Recently, we have developed methods to genetically barcode and quantify muPyVs (36). Here, we use the combination of a non-invasive assay for virus shedding, a library of ∼ 4,000 different barcoded muPyVs, and novel computational approaches to dissect the within-host dynamics of infection using the NGS technology.

Our results demonstrate muPyV as a facile system to study numerous parallel infections in a single host. A large fraction of input barcodes are able to establish long-term infection as shown by studies of shed viruses over time and organ-resident genomes. Different organs display unique compositions of bulk virus genomes, consistent with an inefficient productive exchange of virus between organs. Most importantly, the patterns of barcodes detected in urine differ at early times versus late times of infection. Thus, these multiple and shifting modes of shedding may help explain how small DNA virus populations retain the ability to be maintained and transmitted throughout the life of the host.

## Results

### PyVs tolerate a DNA barcode insertion without overt impact on fitness

We have previously shown that muPyV is amenable to long-term *in vivo* experiments in mice where shedding can be non-invasively measured for the lifespan of the host (34). In a complementary methodological paper, we inserted an 18-nucleotide sequence comprising a barcode of 12 random nucleotides along with a restriction enzyme site into the muPyV genome and showed that barcoded virus stocks gave rise to high titers comparable to the wild-type virus (36). To obtain a quantitative assessment of the 18-base pair insert on viral infection, we conducted viral replication assays and plotted growth curves for the pooled barcoded viruses and compared them to wild-type (no barcode) (Fig. S1A). Although we cannot rule out fitness costs for some individual barcodes, at a broad scale the curves show a similar trend, indicating that numerous different barcodes have little impact on virus fitness.

To determine if a different PyV could also tolerate a barcode insert in the same genomic location, we generated two barcoded BKPyV libraries, which also showed similar final virion concentrations and replication kinetics to wild-type (no barcode) (Fig. S1B and Table S1). We conclude that diverse PyVs can tolerate small inserts between the early and late polyA signals and that barcoded PyVs comprise a DNA virus system to pursue parallel infection studies *in vivo*.

### Longitudinal dynamics of shed viral barcode repertoires indicate smoldering infection from cellular reservoirs

To monitor virus infection, we infected four mice (2 male (“ML”, “MR”) and 2 female (“FL”, “FR”) with 1x 10^6 IU of the barcoded muPyV stock (36) and collected urine approximately 2-3 times per week over a period of 99 days. We note that at day 59 post-infection, one male (“MR”) had to be euthanized due to an abscess that was unrelated to muPyV infection.

qPCR analysis for the presence of viral DNA showed detectable virus genome loads in most urine samples (Fig. 1). For all four mice, we noted a dip in total genome loads beginning at about day 27 post-infection consistent with a transition from acute to persistent infection (22, 34). We noted that while the two female mice displayed a pattern of episodic high shedding events consistent with our previous work using female mice (Fig. 1 and (34)), the two male mice displayed a somewhat different pattern of elevated shedding throughout all time points analyzed. Although this sample size is too small to make definitive conclusions, these observations are at least consistent with the possibility of sex affecting the patterns of PyV shedding. Assaying a small subset of urine samples that displayed high levels of viral DNA from a separate infection showed that all tested samples contained infectious virus, which was not observed in mock-infected controls (Table S2). We conclude that collection of urine samples affords the ability to non-invasively monitor shedding of infectious viruses.

**Figure 1.**
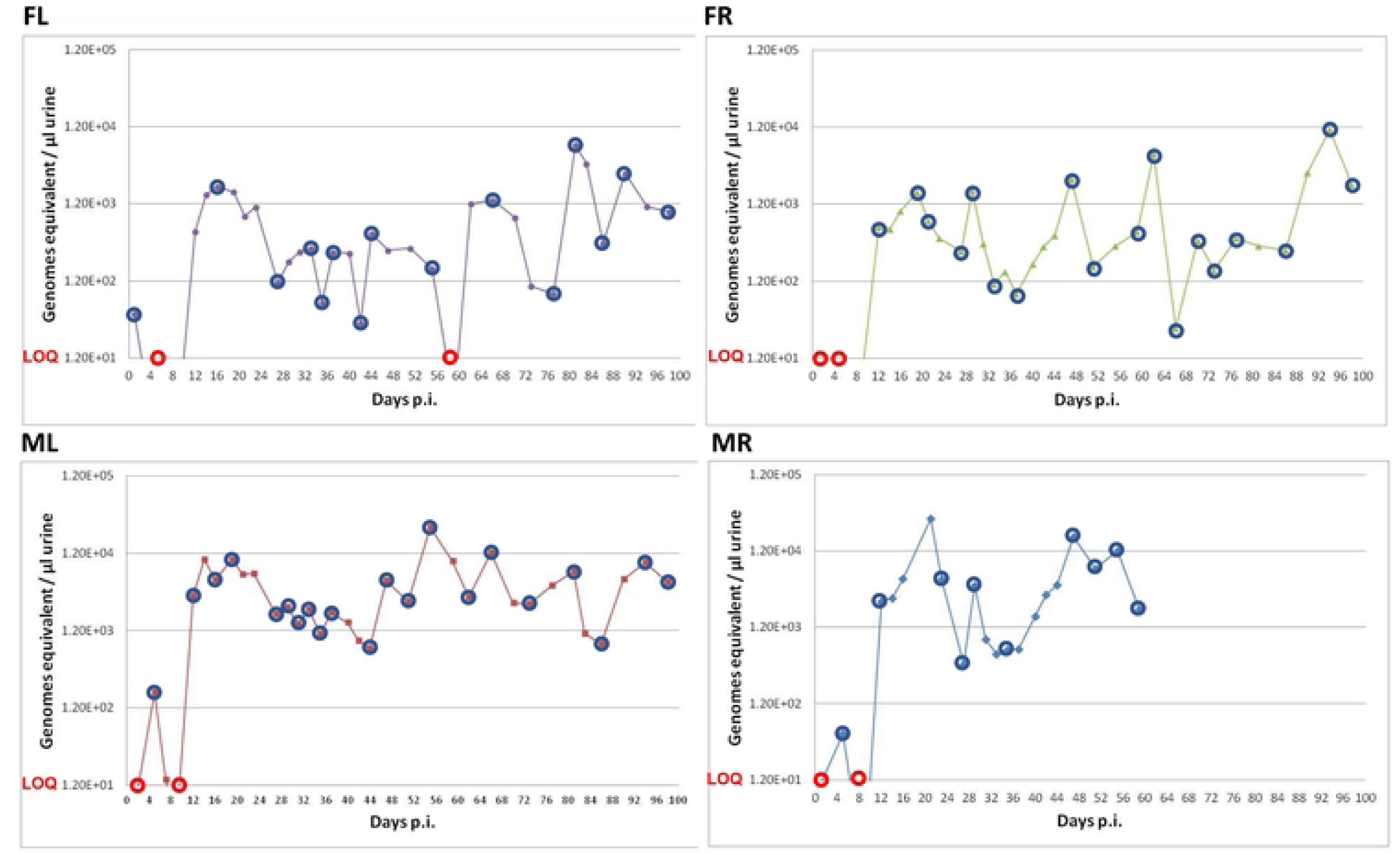
Total amount of muPyV DNA shed in urine over time. Urine was collected from infected mice at multiple times post-infection. muPyV genome equivalents per µl of urine as determined by qPCR are shown. Blue circles indicate urine samples selected to determine the barcode repertoire by Illumina NGS. The few selected samples where viral DNA levels were below the limit of quantification (LOQ) are indicated with red circles.

To determine the composition of barcode repertoires, we generated Illumina NGS libraries for a subset of the urine samples (indicated by red and blue circles in Fig. 1). For every sample, we detected more than 80 unique barcodes, albeit most were of low abundance (Fig. 2). The number of unique barcodes in the samples ranged from approximately 80 to 2000, except for the mouse “FL”, which shed 3132 unique barcodes on the first day post-infection.

**Figure 2.**
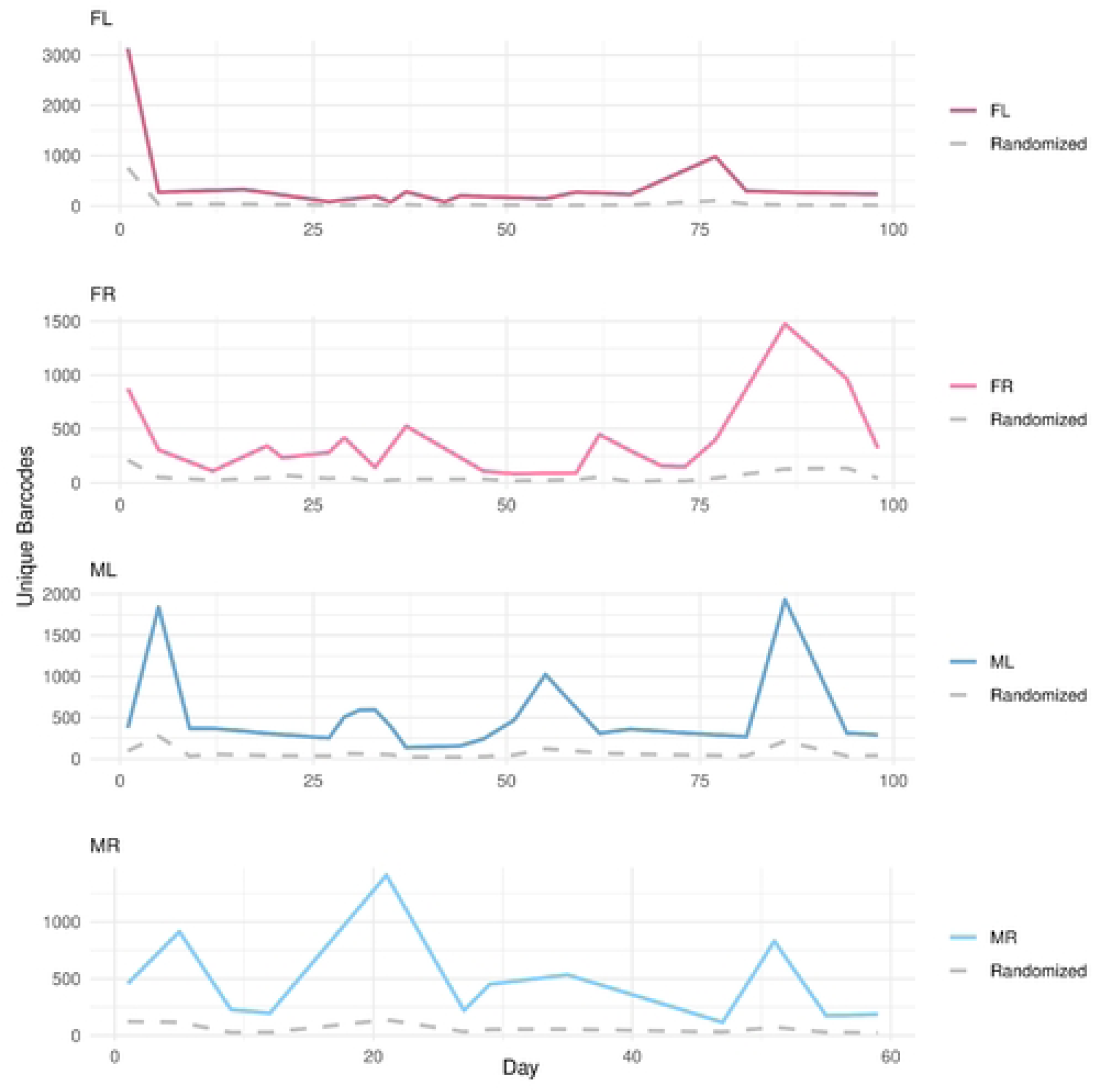
Temporal dynamics of unique barcode detection and enrichment over background. The number of unique barcodes were extracted from Illumina reads for each sample. The number of unique barcodes detected in each mouse over time is plotted. The gray dashed line in each plot represents the *in silico* control, generated by randomly shuffling the nucleotides in each barcode and then associating them with the stock barcodes to measure background noise. The control barcode counts remained stable across all time points, while the number of unique barcodes in the samples exhibited consistent directional trends (upward or downward) over multiple time points.

For urine samples with low virus genome copy numbers, our Illumina analysis was able to detect more genomes than our qPCR analysis (10- 100X more identified via Illumina). To validate the observed pattern of unique barcodes, we performed a control experiment. We randomly shuffled the nucleotides of each barcode in each sample, associated these shuffled barcodes to the stock barcodes using the same method as for the actual samples, and used those mock barcodes as a control. The number of these mock control barcodes, which worked as a measure of background noise, remained relatively low and stable across all time points (Fig. 2). In contrast to the randomized controls, we observed that the number of unique barcodes detected in the actual samples tended to be more abundant and drift downward (down to further down) or upward (up to further up) in a consistent direction across multiple time points. Adjacent time points showing opposite trends (up-to-down or down-to-up) were less commonly observed (Fig. 2). These results are consistent with a smoldering infection, suggesting that individual cellular reservoirs intermittently shed virus over periods spanning at least several days (Fig. 2).

Cosine similarity is a measure of the similarity between two sequences of numbers, where the proportions, rather than the absolute values of the numbers, are compared. In our case the number of reads recalled for an individual barcode are the levels in a sample. Examining cosine similarities between individual time points showed that similar patterns of barcode abundance often clustered together in time (Fig. 3). The exceptions to this were at early times post-inoculation or during later times when there were spikes of highly shed viral DNA. These results suggest that after the early times of infection, low level shedding of muPyV from the same reservoir(s) results in temporally related patterns that can be overshadowed when high shedding events occur.

**Figure 3.**
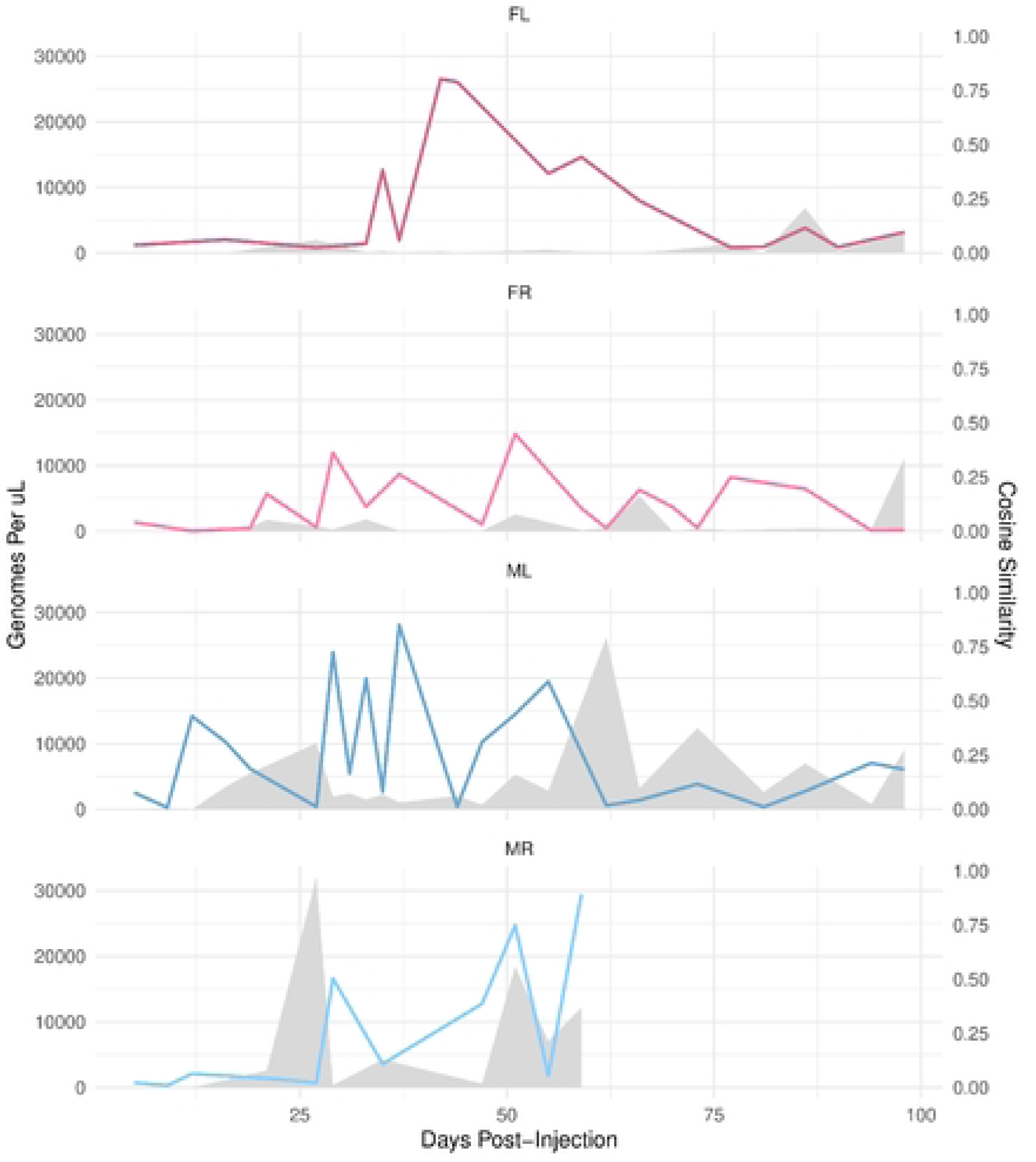
Similarity in patterns of barcodes shed at adjacent times post-infection. Cosine similarity was used to measure the relatedness of patterns between sets of barcodes. The cosine similarity between temporally adjacent time points was plotted to visualize how the relatedness of shed barcode patterns changes over time. Gray area shows the amount of total bulk genomes shed per microliter of urine at each time point. Note that timepoints with abundant total viral DNA shedding generally have lower cosine similarity scores, consistent with less temporal relatedness when large bulk genome shedding events occur.

### A decline in bulk shed barcode diversity occurs early in the acute phase of infection

To understand how the diversity of the barcode repertoire shed in urine evolves over time, we first represented the relative abundance of each barcode (percentage of total) for each sample as donut plots (Fig. 4). Although over 80 different barcodes are detectable at all times, strikingly, for all four mice, we observed a consistent pattern whereby the bulk of viral genomes shed in the early infection was made up of numerous different barcodes which were progressively reduced to only one or a few barcodes accounting for the majority of bulk viral DNA shed during the late times of the persistent infection (most individual barcodes are shown as black or white with the top ten most abundant barcodes shown in color in Fig. 4). Thus, from an initial inoculum of thousands of barcodes, a small number are shed at high levels at late times of persistent infection. The particular barcodes that are shed in each urine sample vary, even within the same animal, but typically the pattern of a few dominant barcodes in any sample tends to persist over several time points.

**Figure 4.**
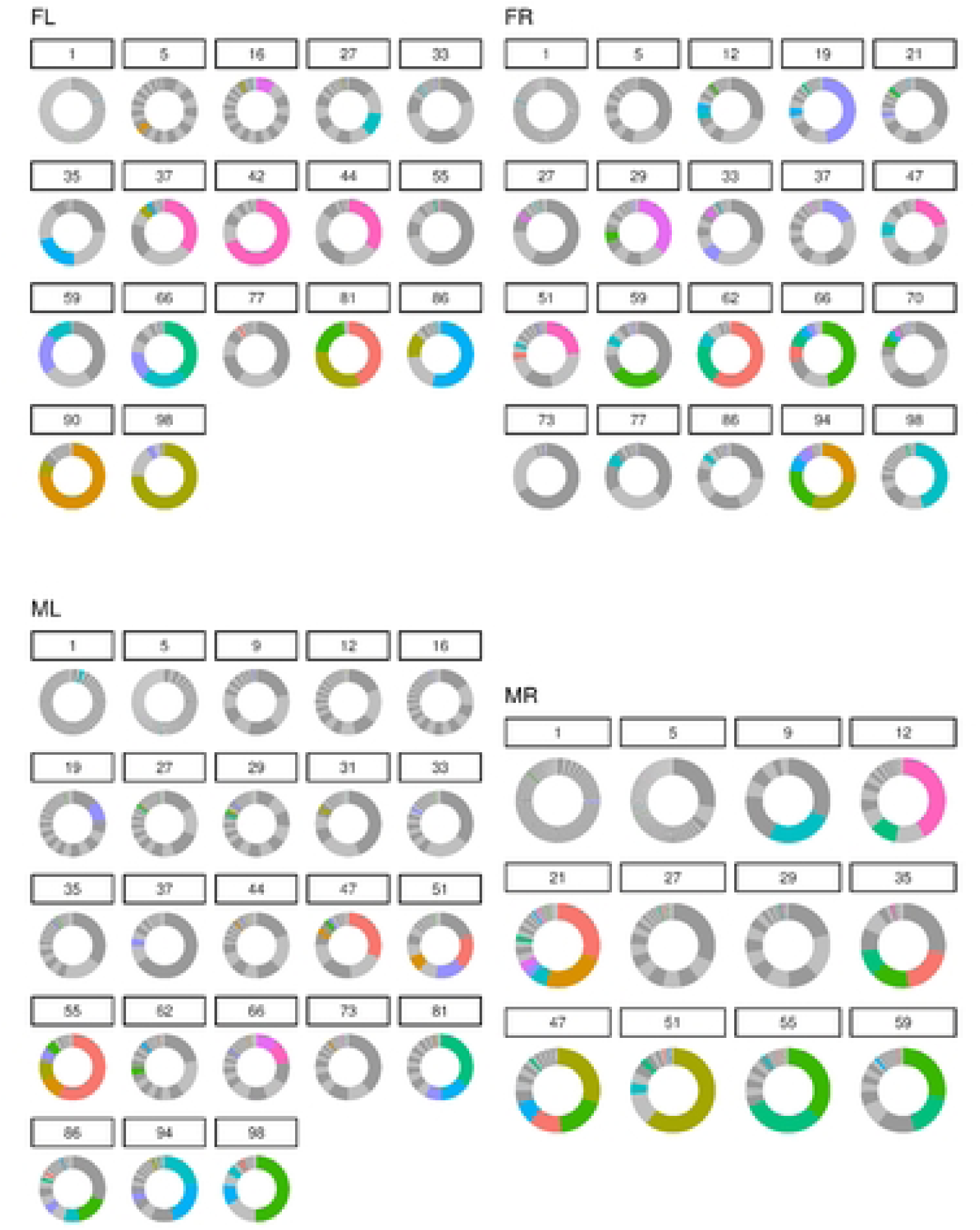
Changes in the diversity of shed barcodes over time. For each time point analyzed, the number of distinct barcodes are plotted on a donut chart with the portion of the circle shaded being proportional to the contribution of that barcode to the overall abundance of bulk barcode DNA present in a sample. Most barcodes are of lower abundance and represented as dark or light gray areas, which take on the appearance of solid gray or alternating stripes. Shading shown in color represents a barcode that was among the top ten most abundant barcodes shed by that mouse over all time points tested. Note, the “9-12 o’clock” region of the plot represents the least abundant barcodes where there are often so many barcodes depicted that the final colors appear a single gray tone. The “12-3 o’clock” region shows the most abundant barcodes for that time point.

As a quantitative measure of diversity, we used Shannon entropy, which captures both the richness and evenness within a population. Using this metric, we observed a steep decline in the diversity of the bulk barcodes shed after the first few days post-inoculation, indicating that the bulk viral population becomes increasingly dominated by a smaller subset of barcodes at later time points (Fig. 5). We observed similar results when we used a third diversity metric, defined as the relative number of different barcodes required to account for 75% of the bulk shed viral DNA at any one time point. Although we cannot rule out that some of the observed barcodes were non-replicating input viruses that were cleared/excreted into urine, these observations are consistent with a high fraction of the input viruses being shed early during infection.

**Figure 5.**
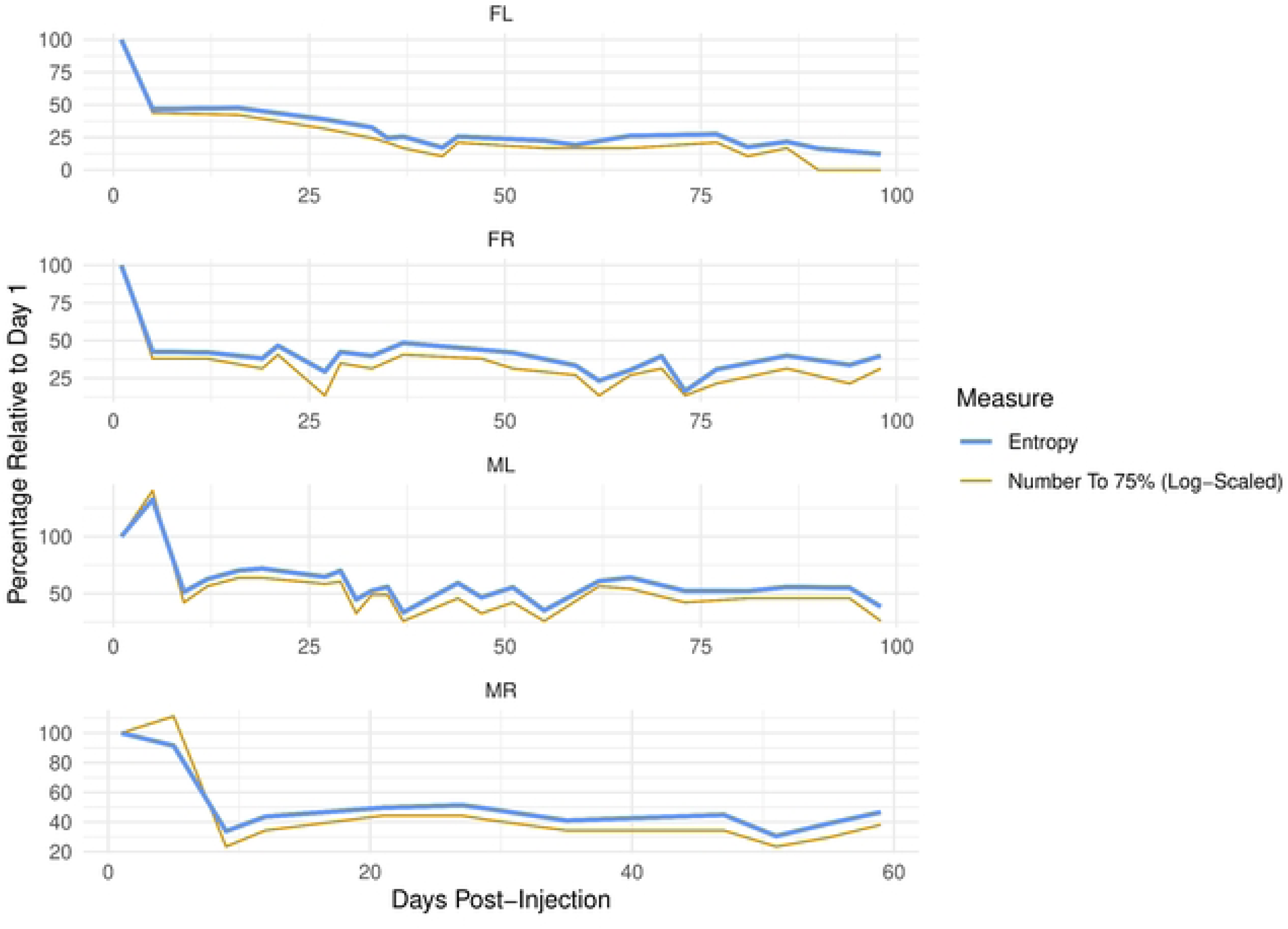
Diversity of barcode repertoire decreases after early times of infection. Shown is the Shannon entropy, which is a measure of complexity (blue line). Overlaid is a plot showing a separate analysis for the number of barcodes it takes to account for 75% of total bulk shed genomes at any single time point (gold). Note, both measures show similar trends of diversity decreasing after the early times post-infection and then stabilizing.

### A small number of barcodes make up the bulk of virus genomes shed during the persistent phase of infection

To further visualize shedding patterns, we plotted the top 30 most abundant individual barcodes shed from each mouse over time (Fig. 6). This analysis showed that, in addition to having at a low shed level of (typically) hundreds of barcodes detectable at all times (as described above), a small number of the barcodes were abundantly shed in punctuated events (Fig. 6). Interestingly, while some of the most abundant barcodes are detectable at different time points post-infection, most barcodes shed at high levels during the late persistent phase of infection were typically detected in only one or a few shedding events (Fig. 6, S2).

**Figure 6.**
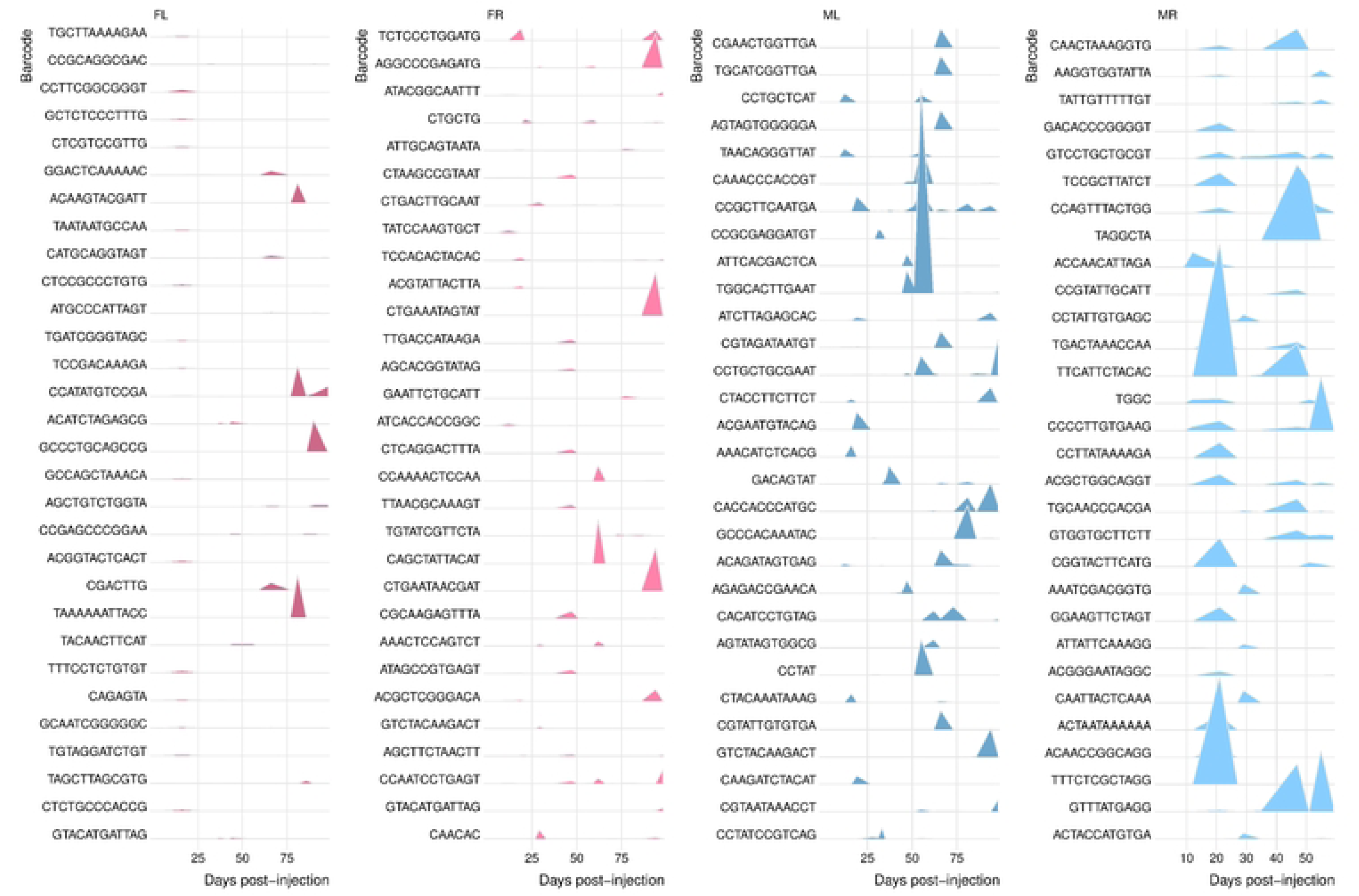
Shedding patterns of the top 30 most abundant individual barcodes found in each mouse during the time course of infection. Linear ridge plots represent genome equivalent copies of the top 30 most abundant barcodes (“most abundant” determined by greatest amount of a barcode shed at any single time point). The height of an individual peak on the vertical axis correlates to the relative linear abundance of each barcode. The horizontal axis corresponds to different times post-infection when levels of shed muPyV DNA in urine was determined.

To comprehend the contribution of the more abundantly shed genomes to total viral genomes shed, we utilized ridge plots to co-visualize the longitudinal shedding of the top 10 most abundant barcodes (“top 10” was determined by the number of reads in any urine sample for an individual mouse). Strikingly, this analysis showed that, except during early times post-inoculation, the top 10 high-level barcodes accounted for the majority of bulk genomes shed in three of the four mice (Fig. 7, S2). The exception to this was mouse “ML”, where the top 10 barcodes still accounted for approximately 40% of total shed barcodes (Fig. 7, S2). These shedding patterns were remarkably consistent across the different host mice. These results are consistent with numerous viruses being able to establish infection and shed continuously at low levels throughout the life of the host, while only a small minority of viruses give rise to subsequent sporadic large shedding events.

**Figure 7.**
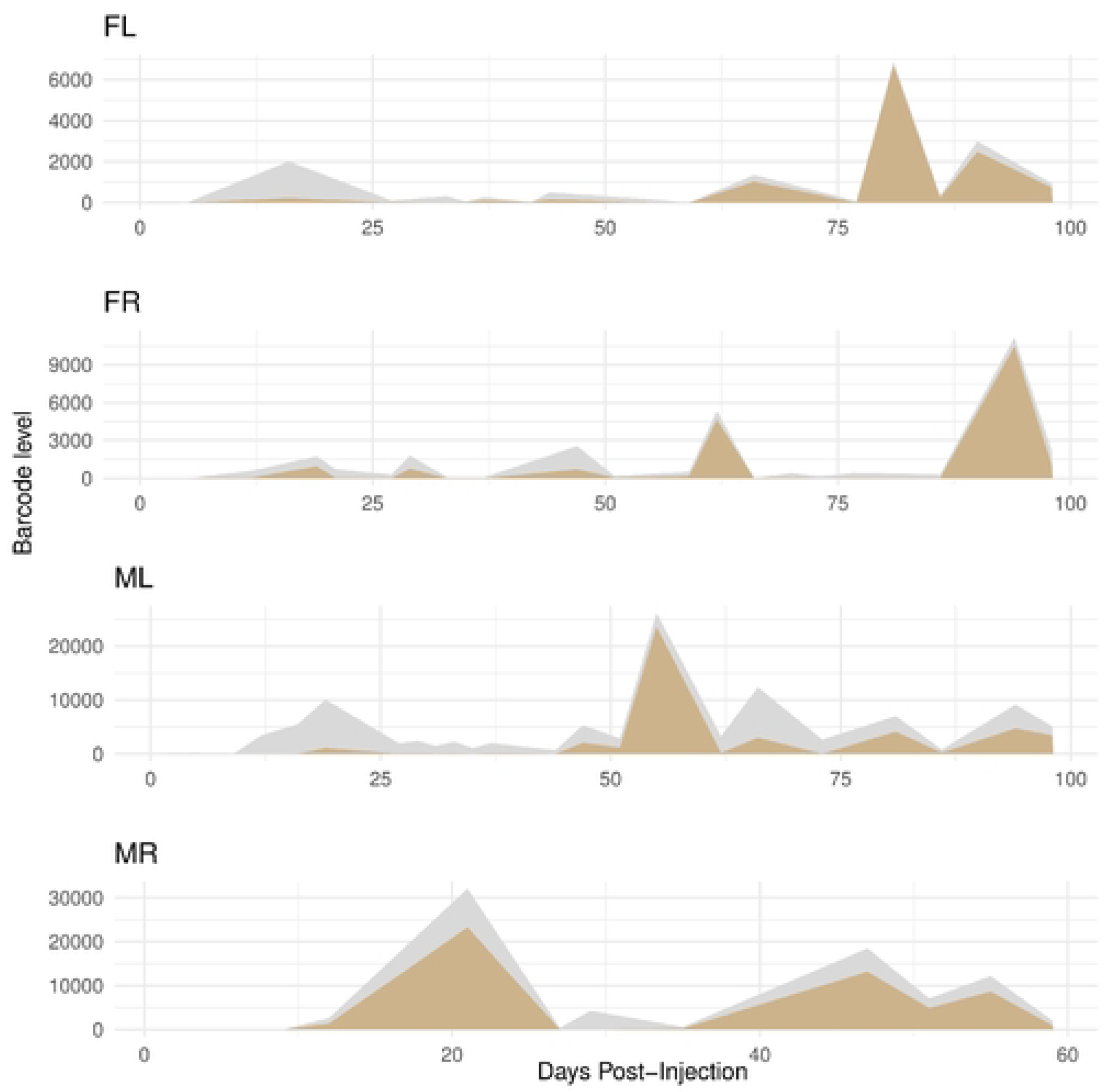
The sum of the 10 most abundantly shed barcodes constitutes the majority of bulk shed muPyV DNA in the late phase of persistent infection. Shown in gold is a ridge plot for each mouse representing the abundance of the sum of the 10 most abundant barcodes shed at each time post-infection (“top 10” determined by the greatest amount of a barcode shed at any single time point). The gray shading represent s the sum total bulk virus genomes shed at any particular time point post-infection. For easy comparison, these plots are duplicated on the bottom of Figure S2.

### Highly shed barcodes were more abundant in the inoculum

We showed previously that barcodes in our muPyV library are not evenly distributed (36). Here, we take advantage of this to determine how the relative input abundance affects the amount of virus genomes shed. Analysis of relative abundance of shed virus genomes in urine showed that the 40 barcodes combined from the top 10 most abundant barcodes from each of the 4 mice had a median rank of 456 - 1266.5 out of 4012 total in our input library (i.e., a median rank range in the top 11.36 % - 31.56% most abundant of the input barcodes) (Table S3). These results demonstrate that genomes abundantly shed at later times of infection were overrepresented in the input inoculum. This suggests that the initial viral load influences the mode of infection and transmission of the virus. However, factors other than the input abundance must also contribute since a minority of the more abundantly shed genomes were ranked in the bottom half or quartile of abundance of the input inoculum library (Fig. 8). Analysis of the barcode GC content and barcode length didn’t show any trend of enrichment over the course of the experiment (Fig. S3, S4), suggesting that general sequence features of the barcode do not confer a selective advantage leading to the highly shed barcodes in our system. Thus, input abundance and other factors influence the propensity for organ persistence and shedding in urine.

**Figure 8.**
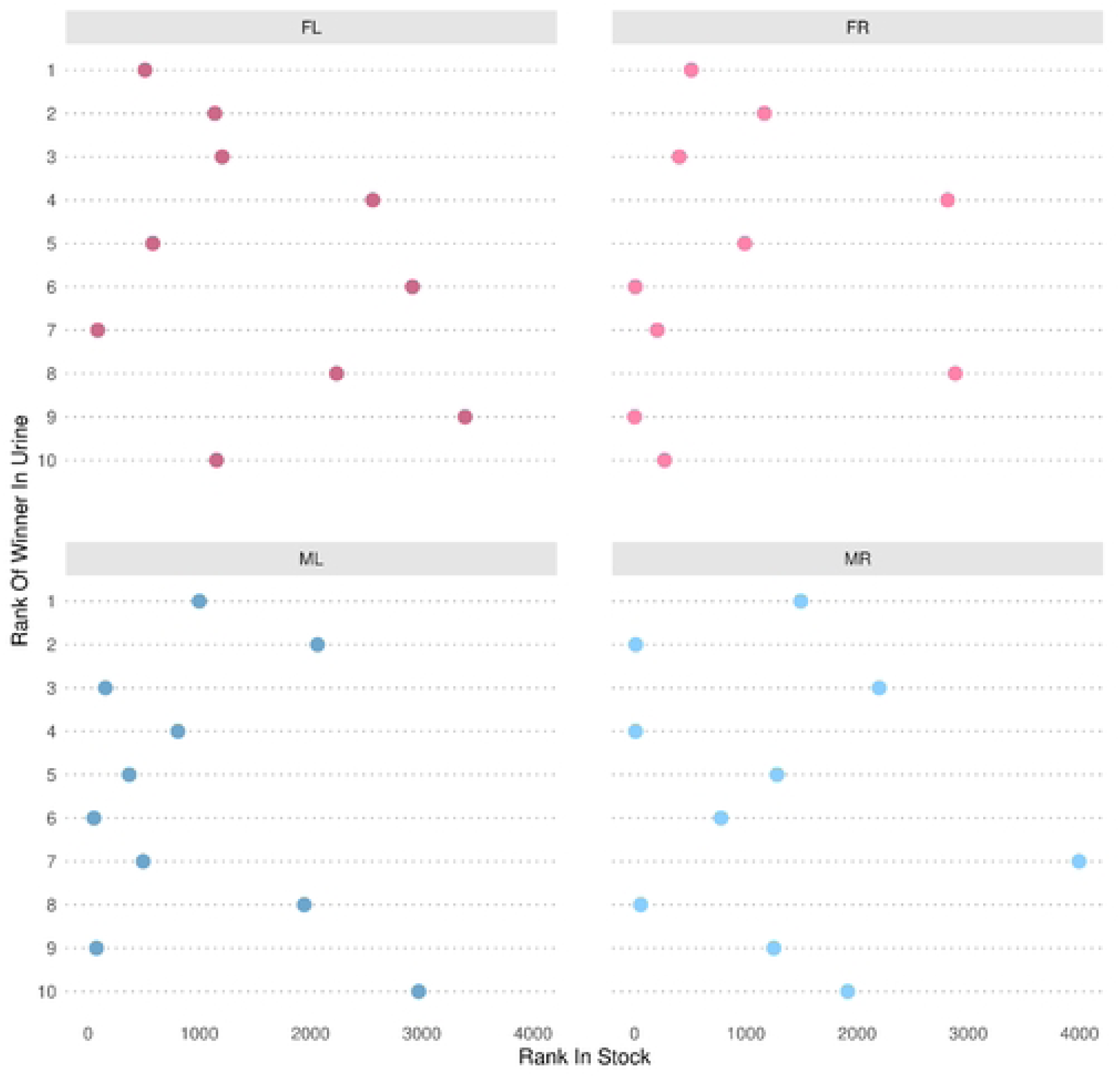
Rank of the most abundant shed barcodes in the inoculum virus stock. Shown is the rank in the virus stock of the top 10 most abundant barcodes shed in urine (“top 10” determined by greatest amount of a barcode shed at any single time point). A lower rank (more towards the left side) indicates higher abundance in the virus stock. Note, the general trend shows that many, but not all, of the most abundant shed barcodes were among those relatively more abundant in the initial inoculum virus stock.

### Individual organs possess unique patterns of relative barcode abundance

At the conclusion of our longitudinal study (99 d.p.i. for three mice, 59 d.p.i. for one mouse), we harvested various tissues and organs, quantified the genome equivalent copy number by qPCR (Fig. S5), generated Illumina NextSeq NGS libraries, and quantified individual barcoded genome equivalents present in each tissue/organ type. Consistent with previous reports, readily detectable viral DNA was observed in diverse tissue types (Fig. S5) (34, 37). In the combined tissues analyzed for each mouse, we observed that 92-96% barcodes from the input library were represented, except for the mouse “FL”. Even in the case of the mouse “FL”, the representation was still high at ∼73%. By Fisher’s exact test, we found that all mice showed a significant overlap between the top 5% of urine and tissue barcodes, consistent with the propensity to be shed in urine correlating with enhanced residence in tissues (Fig. S6). Although there was substantial variation in the relative levels of individual barcodes, the vast majority of barcodes from the inoculum established and maintained infections in the combined organs of each mouse.

Similar to what we observed for the most abundant viral genomes shed in urine, no trend of GC content enrichment was detected in the barcode sequence of the most abundant genomes in organs (Fig. S7). Moreover, the most abundant barcodes in organs were overrepresented in the input virus library (median rank range of 292.5– 1312.5 out of 4012 total input barcodes) although these barcodes were not necessarily the same as those enriched in urine (Table S3). Like in urine, factors other than the input abundance also contribute to the abundance in organs since a minority of the more abundant genomes in organs were ranked in the bottom half or quartile of abundance of the input inoculum library (Fig. 9). Although some of the relatively more prevalent barcodes in one organ were also prevalent in other tissues within an individual mouse, the pattern of barcodes with the highest relative abundance was distinct in each tissue type for any single mouse as shown by low correlation of barcode relative abundance between organs (Fig. 10). These data suggest that enhanced levels of a particular virus in the inoculum increase the likelihood of higher relative abundance in organs, but there is limited productive exchange of abundant genomes between organs.

**Figure 9.**
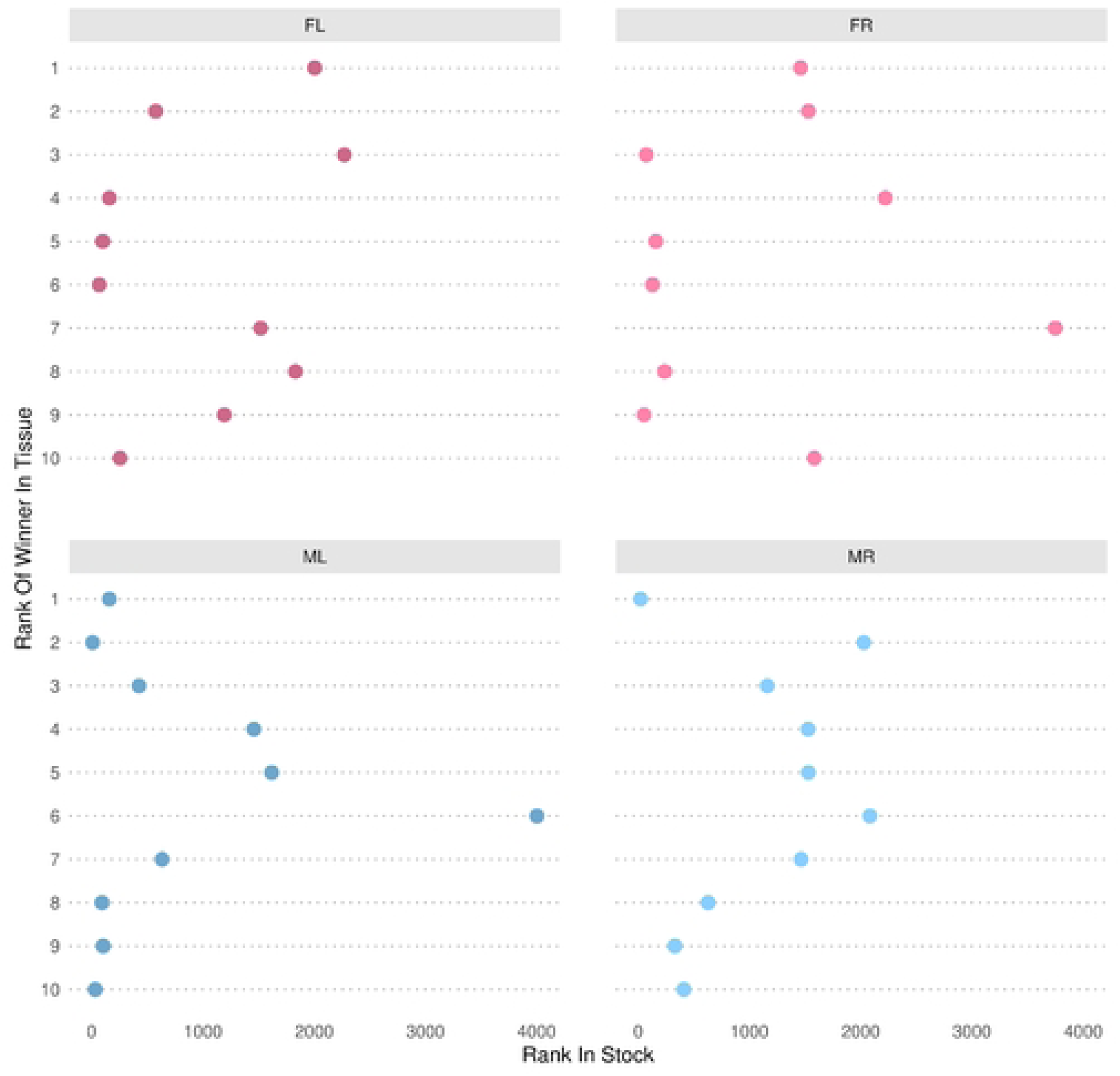
Rank of the most abundant barcodes in assayed mouse tissues in the inoculum stock. Shown is the rank in the virus inoculum stock of the top 10 most abundant barcodes (“top 10” determined by the greatest amount of a barcode in any tissue for an individual mouse). A lower rank (more towards the left side) indicates higher abundance in the virus stock. Note, similar to what is detected shed in urine, the general trend shows that many, but not all, of the most abundant shed barcodes were among those relatively more abundant in the initial inoculum virus stock.

**Figure 10.**
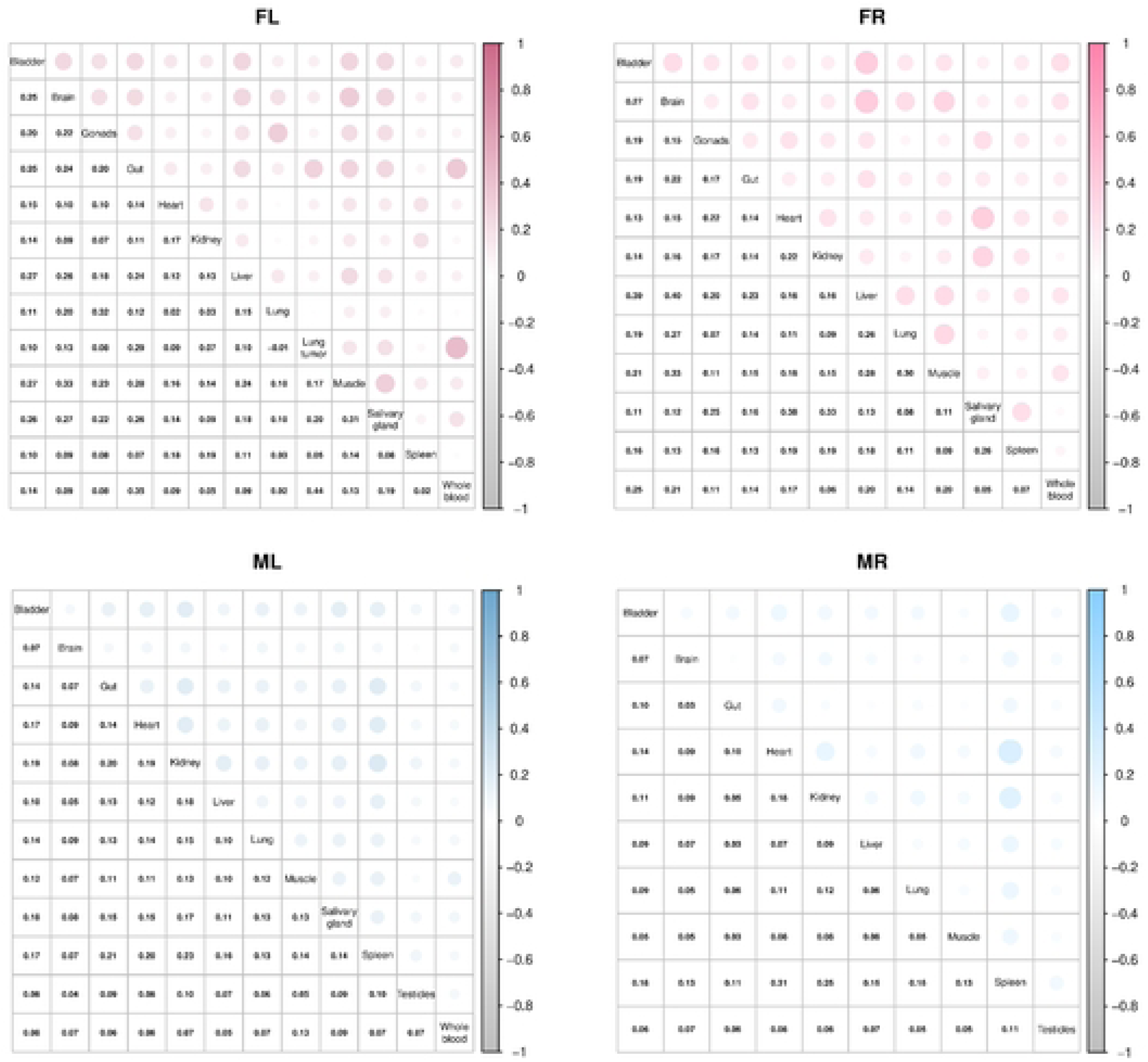
Low correlation of barcodes repertoires amongst different tissues. Shown is the Spearman correlation coefficient of relative barcode abundance between tissues in each mouse. The top right corner represents the correlation by the size of circle and intensity of the shading between any two organs; the bottom left corner shows the numerical value of Spearman correlation coefficient between different pairs of organs withing an individual mouse. These data demonstrate that tissues have unique repertoires of virus genomes, consistent with limited exchange of virus genomes between tissues.

### High levels of viral genomes in tissues do not necessarily give rise to infectious viruses

After euthanasia, we noticed that a tumor spontaneously developed in the lung of one of the mice (FL). Interestingly, this tumor contained a high-level of muPyV DNA (2.42E+08 genomes equivalents per µg of total DNA). Analysis of Illumina reads showed that, although non-tumor lung tissue also contained a high-level of muPyV genomes, the tumor predominantly contained a single barcode. To understand the reason of high levels of muPyV DNA in this tumor and lung, we sequenced the full muPyV genomes by Sanger sequencing. The sample from the tumor showed a deletion of 1052 bp, starting from the Non-Coding Control Region (NCCR) and ending in frame in the late region coding for the capsid protein VP1 (with complete deletion of VP2 and VP3 regions) with no other mutations detected (Table S4). Interestingly, the sequencing of two muPyV genomes from the lung of the same female showed a smaller deletion (625 bp) in the same genomic region ending in the VP2 and VP3 coding region, along with mutations in the NCCR of the two sequenced genomes and additional mutations in the T antigen coding region of one of the two genomes. In the lung of the other female mouse, we also observed a similar deletion (1504 bp) with no additional mutations (Table S4).

These large deletions likely prevent the expression of functional capsid proteins, thus preventing production of virions and cell lysis. Such high copy number of muPyV genomes would also be expected to be able to drive the expression of high-levels of early genes that are known to exert a mitogenic effect (38–40). Inverted PCR amplification showed a product of expected length for a full genome, which was confirmed by Sanger sequencing, suggesting that these genomes are circular and non-integrated within the host genome. These results suggest that tumorigenesis in this instance occurred via mechanisms consistent with episomal-muPyV-driven tumors (41). These findings also suggest one mechanism whereby neighboring organs and tumors can harbor high levels of viral genomes without giving rise to infectious viruses capable of seeding new infections in neighboring tissues.

To determine which organs contribute to the shed virus detected in urine, we compared the relative ranks of barcode abundance in the various organs to their rank in urine collected on the last day before the mice were sacrificed. This analysis revealed a strong signature of shared enrichment in the kidneys of 3 of the 4 mice (Fig. S8). Notably, no other organs displayed a similar strong shared enrichment. These findings are consistent with the kidney being a primary site of production for virus shed in the urine, as previously established for muPyV and other PyVs (26, 35, 42, 43). These observations further emphasize the apparent siloed nature of virus genomes between different organs.

## Discussion

Members of some DNA virus families can undergo life-long persistent infections (14, 15). Herpesviruses have large genomes with numerous genes to manipulate the host response to infection, as well as defined mechanisms of latent and lytic infections (44–59). In contrast, small DNA viruses like polyomaviruses, can also undergo long-term and sometimes lifelong persistent infections with substantially smaller genetic toolsets and little understanding of the mechanisms involved (25). Here, we unveil basic aspects of acute and persistent infection of a prototypic model PyV. Our work shows that thousands of muPyVs can initiate infection, disseminate to multiple organs/tissue types, persist for months after infection in diverse organs, and most impressively, shed essentially continuously via temporally shifting patterns of virus release. What emerges is a model whereby PyVs have evolved multiple and overlapping strategies for transmission.

We utilized a genetic barcode approach whereby we tagged a region of muPyV genome with thousands of random barcodes to study patterns of infection and shedding (36). Similar to some human PyVs (26, 60, 61), muPyV sheds infectious virus in urine (34, 35), which enabled us to non-invasively probe longitudinal infection dynamics of a single host infected with numerous viruses simultaneously. Our results show that higher relative barcode levels in the inoculum increased the likelihood of greater abundance being shed in urine months after initial infection. Similar trends are observed in organs during late persistence. However, additional unknown factors also contribute to the chances of being detected in organs and being shed in urine as a minority of these most abundant virus barcodes were also among the relatively less abundant in the inoculum.

Interestingly, distinct populations of relative barcode abundance are detected in individual tissues at late times of persistence. Only the kidney appeared to contribute to viruses shed in urine underscoring a lack of virus genome exchange between organs. In the tumor that arose in the lung of one mouse, we observed a dominant abundant muPyV genome. This genome was non-integrated and contained a large deletion ablating most late gene expression. If a large fraction of genomes are maintained as episomes in normal tissues, this could explain how barcodes drift in abundance specific to individual organs/tissues. It would be interesting in future studies to determine the fraction of organ-resident genomes that are episomal and how these are maintained. Irrespective of the mechanism, these findings demonstrate that PyV infections can be established and persist in individual organs with little productive exchange of abundant genomes between neighboring organs.

Remarkably, throughout essentially all times post-infection up to day 99 post-inoculation, urine contained many (> 80) detectable low abundance barcodes. However, our analysis reveals a noticeable shift in the pattern of shed barcodes in the early acute versus late persistent phases. At early times post infection (before day 6), a high diversity of barcodes comprise the bulk shed genomes at any one time. This is consistent with numerous reservoirs of infection each shedding virus. But, at around 6 days post-infection, there is a shift to a lower diversity of the bulk shed genomes. During the persistent phase, despite thousands of barcodes still being detectable at low levels, large amounts of a few barcodes tend to dominate the bulk of total shed genomes. Highly abundant shed barcodes occurred at punctuated times with only a few abundant (typically less than 10) barcodes comprising the bulk mass of shed genomes at any one time. These results suggest that some viruses emerge from a limited reservoir of infected cells that are able to give rise to a large amount of shed virus.

One hypothesis to explain the punctuated highly shed genomes is something akin to herpesvirus lytic reaction. However, other models are possible and proving that PyVs can undergo true latent/lytic infections akin to herpesviruses would require studies demonstrating alternative and reversible viral gene expression programs (45). We note that both smoldering and latent/lytic infectious cycle have previously been proposed for PyVs (21, 25, 62–64). Interestingly, type I interferon (IFN) has been shown to promote Herpes Simplex I latency (65) and can foster persistent infection of other diverse viruses (66). IFN has also been linked to altering infection of diverse types of PyVs (67–69) and has been shown to associate with BKPyV and JCPyV persistent infection in select cultured primary cells (70, 71). Thus, it would be interesting to probe a role for IFN in modulating muPyV shedding patterns.

A major question is what biological cues drive the occasional punctuated highly shed muPyV genomes. In addition to the innate IFN response as discussed above, two additional areas to explore include the adaptive immune response and sex hormones, both of which have been shown to affect PyV infection (72–74). Although our sample sizes are too small to make conclusions regarding sex differences, it is intriguing to note the apparent different patterns of shedding observed between male and female mice (Fig. 1). For all mice we examined, the consistency of punctuated release of large amounts of virus we detect is similar to what is observed in humans naturally infected with BKPyV (75, 76). Similarly, at least some immunocompetent humans appear to continuously shed low levels of BKPyV (61, 77). These observations support that the muPyV model recapitulates the biology of diverse urinary-tract-tropic PyVs. Although rare, serious and sometimes fatal disease can arise from the emergence from persistence of PyVs in immunocompromised hosts (25, 72). Determining the biological factors that shift to a less diverse pool of shed viruses and what controls reactivation of the minority of abundantly shed genomes in muPyV may lead to new strategies for controlling PyV infections in the clinic.

In conclusion, we demonstrate that thousands of different muPyV genomes are able to establish infection in diverse organs. Although numerous genomes persist in organs and are continuously shed in urine, the diversity of the bulk genomes shed decreases days after inoculation. Limited productive exchange of abundant barcodes is observed between organs consistent with our observation that most organs are not contributing to the barcodes that are eventually shed in urine. Only an atypical minority of barcodes are shed at high levels late during persistent infection that may be responsible for the dissemination of the virus to new hosts. Together, the existence of multiple and shifting patterns of PyV shedding that span the early acute to late persistent phases may help to explain the ubiquity and success of these common evolutionary co-passengers. Most importantly, the identification of the host factors responsible for the virus reactivation during the persistent infection may lead to the development of new therapeutic options against life-threatening diseases in humans.

## Limitations of this study

It remains unknown what fraction of the shed barcodes detected are due to replicating virus versus deriving from the initial non-replicated inoculum that may end up passing through the urinary tract. This is important to note in light of the decrease in diversity of barcodes that we detect after the early times post-infection. Our qPCR analysis is sufficient to identify trends in bulk shedding across timepoints but these values cannot be considered absolute at low copy numbers because our Illumina analysis suggests that our qPCR underestimates the true genome copy number. Some samples, including some organs and early times post-inoculation urine, had substantially less total viral material than other samples and this potentially could affect the accuracy of the barcode composition. Although the two female mice studied here shed bulk viral DNA in a pattern consistent with our previous work conducted with female mice (34), the two male mice appear to have a different pattern with less periodicity and retaining higher shedding. Although provocative and consistent with potential sex differences in shedding, with such a small number of mice studied, it remains possible that these observations of apparent male-specific shedding patterns were detected by random chance. It would be interesting to follow up with additional studies on possible sex differences of PyV shedding. Our observations of different patterns of shedding, with spikes of a small number of highly shed barcodes overlaid on a background of constant low-level shedding of a large number of barcodes, are consistent with a population of cells undergoing low-level smoldering infection with possible reactivation of high levels of viral replication in only subsets of infected cells. Such a model of multiple modes of shedding may help explain successful dissemination of PyVs.

However, our study did not directly probe transmission. Further, although viral DNA in urine associates with infectious virus, this association is not linear and was not determined specifically from urine samples with spikes of highly shed barcoded virus (Table S2). Therefore, it remains unclear how much the punctuated high viral DNA shedding events matter for the fitness of transmission. Finally, it is interesting that only a small subset of barcodes are highly shed at any one time point during the late stages of persistent infection. This observation is consistent with one or small number of reservoirs receiving signal to reactivate high levels of virus replication and shed large amounts of virus. However, little is known about the cells that comprise the reservoirs where these viruses emerge from including how many reservoirs are shedding at any one time, how many cells make up such a reservoir(s), and whether there are systemic cues signaling one or multiple cells or reservoirs to shed highly at the same time. Future work is required to resolve these questions but elucidating the host pathways involved in these cues has potential for understanding important basic and clinically relevant aspects of PyV biology.

## Acknowledgements

We dedicate this paper to the memory of Benni Goetz, who worked tirelessly to complete this project and whose unique intellect and creativity permeate much of this work. We thank James J. Bull, University of Idaho, for helpful conversations regarding experimental design, controls, and applications of this work, and for helping to shape some of the arguments put forth in this manuscript. We thank Justin Lau, Upasana Nepal and Aaron Stark for technical assistance maintaining mice and collecting urine samples. We thank Sullivan Lab members past and present for suggestions to visualize data, technical advice, and proofing the manuscript.

## Methods

### Cells

The NMuMG cells (ATCC^®^, # CRL-1636) were kindly provided by Prof. Aron Lukacher (Pennsylvania State University Medical School at Hershey) and maintained in DMEM supplemented with 10% (v/v) fetal bovine serum and 1% (v/v) penicillin-streptomycin (78). The RPTE cells were purchased from Lonza (catalog # CC-2553) and maintained at very low passage in REGM^™^ Renal Epithelial Cell Growth Medium BulletKit^™^ (Lonza) at 37°C with 5% CO2, as recommended.

## Viruses

### Construction of the barcoded muPyVs library and generation of the muPyV stocks

The methods used to generate and determine the concentration of the muPyV wild-type stock and the barcoded muPyVs library have been described previously (36). In brief, barcoded muPyVs have been engineered to carry a barcode made of 12 random bases and a restriction site (MfeI) in between the two polyadenylation signals. Wild-type and barcoded viruses have been generated by excision of viral genomes from the bacterial plasmid, re-circularized, and transfected into NMuMG cells. Virus stocks were harvested, propagated, and tittered by immunofluorescence assay as previously described (36).

### Construction of the barcoded BKPyV libraries and generation of the BKPyV stocks

Two barcoded BKPyV libraries, namely BKBC1 and BKBC2, were constructed based on the position of the barcode between the two poly A signals. The BKPyV Dunlop strain (GenBank accession No KP412983) was used as a background for the construction of the two BKPyV libraries. pUC19 vector containing the full genome of BKPyV Dunlop at the BamHI site (kindly provided by Walter Atwood, Brown University) was amplified by reverse PCR using the Phusion high-fidelity polymerase (NEB) and the two following pairs of primers previously 5’ phosphorylated, BK_BCF1 5’-NNNNNNCAATTGAATAAATGCTGCTTTTGTATAAGCCA-3’ with BK_BCR1 5’- NNNNNNAAATGTATATGTACAATAAAAGCACC-3’ for the BKBC1 construct and BK_BCF2 5’- NNNNNNGTACATATACATTTAATAAATGCTGC-3’ with BK_BCR2 5’-NNNNNNCAATTGAATAAAAGCACCTGTTTAAAGCAT-3’ for BKBC2. The PCR products were digested by DpnI (NEB) (79) and gel purified before the self-ligation step using the T4 DNA ligase (NEB). Then, the ligation products were purified and used to transform MAX Efficiency^®^ DH5α^™^ competent cells (Invitrogen). Transformed bacteria were cultured overnight in LB medium + 100μg/mL of ampicillin and the pool of plasmids was purified. In parallel, the number of transformed bacteria was evaluated on LB+100μg/mL of ampicillin plates. We estimated that the pool contains 4842 and 5274 colonies for BKBC1 and BKBC2, respectively. The presence of the barcode was directly confirmed by Sanger sequencing on 10 colonies for each construct using the BKFull3 primer 5’-TCCCAGGTAATGAATACTGAC-3’. Once the presence of the barcode was confirmed, the two pools of plasmids containing the barcoded BKPyV genome as well as the BKPyV genome without barcode (wild-type) were digested by BamHI-HF (NEB) to separate the virus genomes from the vector by gel purification. Then, the BKPyV genomes were self-ligated overnight at 16°C using the T4 DNA ligase (NEB) and purified prior to transfection into 293TT cells using the Lipofectamin2000 reagent (Invitrogen). The cells and the supernatants were collected at 90% CPE (7 days post-transfection), subjected to freeze/thaw cycles, and cleared by centrifugation at 4°C. Fresh 293TT cells were infected with the crude lysate at 37°C and cultured until 80% CPE (day 14 post-infection). The cells and the supernatants were harvested, subjected to freeze/thaw cycles, and cleared by centrifugation. Fresh RPTEC cells were infected with cleared BKPyV crude lysates at M.O.I. 0.01 and supernatants were collected at 80-90% CPE (day 13 post-infection). Supernatants were cleared by centrifugation at 6,000 rpm for 40 min at 4°C and layered on a 20% sucrose cushion prepared in buffer A (10mM HEPES, pH8; 1mM CaCl2; 1 mM MgCl2; 5mM KCl) and centrifuged at 25,000 rpm for 3 hours at 4°C. BKPyV stocks were resuspended in buffer A, centrifuged at 13,000 rpm for 5 min at 4°C, aliquoted and stored at -80°C until use.

To confirm the presence of the barcode in the virus libraries, an aliquot of the virus stocks was DNAse I treated before the DNA purification step using the QIAamp^®^ DNA Mini Kit (Qiagen) and the region of the virus genomes surrounding the poly- A signals was amplified by PCR with Taq DNA polymerase (NEB) and the primers pair BKBCenrichF 5′-GGTTAGGGTGTTTGATGGCA-3′ and BKBCSeqR2 5′- CCCCTGCTGAAGATTCCCAA-3′. The PCR products were purified, cloned with TOPO^®^ TA Cloning kit (Invitrogen) and Sanger sequenced. To confirm the NCCR region integrity, the virus stocks DNA was amplified using NCCRF1 5′-ATTTCCCCAGGCAGCTCTTT-3′ and BKBCSeqR2 5′-CCGTCTACACTGTCTTCACCT-3′, cloned with TOPO^®^ TA Cloning kit (Invitrogen) and Sanger sequenced.

### BKPyV titration by immunofluorescence assay

The RPTE cells were infected in duplicate with serial dilutions of the virus sample for 1 hour at 37°C. 48 hours p.i. cells were successively incubated at room temperature with PBS containing 4% paraformaldehyde for 20 min, 0.1% Triton X-100 for 5 min, 1% goat serum for 1 hour, BKPyV VP1 monoclonal antibody (M19), clone 5E6 (Abnova) diluted 1:400 for 1 hour and stained with Alexa Fluor 488 goat anti-mouse IgG (ThermoFisher Scientific) for 1 hour. The number of stained cells per field was counted under an inverted fluorescence microscope (Leica), and infectious titers (IU/ml) were calculated.

### Growth curve of the barcoded BKPyVs libraries

The RPTEC cells were infected with the barcoded BKPyVs libraries or the BKPyV Dunlop wild-type stock at M.O.I. 1 for 1 hour. Then, the supernatant was removed, and unabsorbed viruses were washed out. Infected cells were cultured for 48, 72, 96, and 120 h p.i. At each time point, the supernatants were collected and cleared by centrifugation before titration.

### Growth curve of the barcoded muPyVs library

The NMuMG cells were infected with the barcoded muPyVs library or the muPyV PTA wild-type virus stock at M.O.I. 5 for 1 hour at 37°C. Then, the supernatant was removed, and unabsorbed viruses were washed out. Infected cells were cultured for 4, 20, 24, 28, 40, and 48 h p.i. At each time point, the supernatant was discarded and cells were washed and resuspended in PBS. Cell pellets were freeze/thawed and the supernatants were collected by centrifugation before titration. To visualize and quantify infected cells, immunofluorescence microscopy was conducted similarly to as described above for BKPyV, except anti-muPyV VP1 rabbit polyclonal antibody (a gift from Richard Consigi) was used.

### Barcoded virus longitudinal shedding, infections and sample collection in mice

All animal procedures were performed in compliance with the approved University of Texas at Austin Animal Care and Use Committee protocol and were subject to veterinary approval. Two male and two female FVB/NJ mice, between 9 and 10 weeks of age were inoculated with 10^6^ IU of the barcoded muPyVs via intraperitoneal (i.p.) injection. To collect urine, mice were placed on a separated microisolator cage with a wire-bottom insert and lined with plastic wrap for 2 to 4 hours. Urine samples were stored at -80°C until subsequent analysis. At the end of the longitudinal study (i.e. 59 days post-infection (d.p.i.) for 1 male and 99 d.p.i. for the other three mice), the bladder, brain, gut, heart, kidney, liver, lung, muscle, salivary gland, spleen, testicles, whole blood and 1 lung tumor were harvested and split in 2 aliquots for DNA purification. Samples for DNA purification were snap- frozen in liquid nitrogen and stored at -80°C until use.

### DNA extraction for samples analyzed in the longitudinal study

DNA was purified from 50μl of urine (or less when low volumes were collected) using the QIAamp viral RNA mini kit (Qiagen), following the manufacturer’s protocol. DNA was eluted in 60 μl of Tris-EDTA buffer and stored at -20°C until use. For qPCR assays, 2 μl of eluted DNA was assayed whereas typically 20 μl of eluted DNA was used as input for Illumina reactions.

DNA from 25mg of organs (or less if the sample size was not enough and only 10mg of spleen) was extracted using a bead beater (Bead Mill4, Fisher Scientific) and the QIAamp Fast DNA Tissue kit (Qiagen), following the manufacturer’s protocol. DNA was purified from 140μl of whole blood with the QIAamp DNA Mini kit (Qiagen) following the manufacturer’s protocol. An additional RNase A digestion step was performed for the whole blood samples. DNA was eluted in 200μl of AE buffer. The DNA concentration was measured by the NanoDrop spectrophotometer and the integrity of each sample was checked on a 0.8% agarose gel. Samples were stored at -20°C until use.

### Real time-qPCR assay for the muPyV DNA quantification

The 20μl qPCR mixture contained 10μl of PerfeCTa^®^ SYBR^®^ Green FastMix^®^, ROX^™^ (Quantabio), 8pmol of each primer PTA/PTA-dl1013 sense 5’-GATGAGCTGGGGTACTTGT-3’ and PTA/PTA-dl1013 antisense 5’-TGTATCCAGAAAGCGACCAAG-3’ and 2μl of eluted DNA solution as template per reaction. qPCR reactions consisted of an initial denaturation step of 10 min at 95°C, followed by 40 cycles of 15 sec at 95°C, 30 sec at 60°C. Fluorescence was measured during each extension step. The specificity of each PCR was checked by melting curve analysis. Real time-qPCR was performed on a StepOnePlus real-time PCR system (Applied Biosystems) and analyzed using the StepOne software v2.2.2. The copy number of muPyV DNA was determined via reference to a standard curve also prepared in duplicate by ten-fold serial dilution of a single pBluescript-sk+PTA barcoded vector. DNA from tissues was freshly diluted to 100ng/μl or 10ng/μl in water just before qPCR. The copy number of muPyV DNA was normalized per μl of samples or per μg of total DNA for tissues. The PCR reaction was performed in duplicate with the average of two replicates comprising the final quantification. The limit of detection was 10 copies per reaction.

### *In vivo* infection of polyomavirus and urine collection to detect infectious virus in urine

The mice used were C57BL/6N (Taconic) background. To confirm highly positive samples contained infectious virus, six C57BL/6N mice were initially infected with 1x10E+06 IU/mouse of the barcoded muPyVs library and the wild-type PTA muPyV stock in a 1:1 ratio by intraperitoneal inoculation. Four mock-infected mice were also included as negative controls. Urine was collected by placing each mouse in a microisolator cage with a wire-bottom insert and lined with plastic wrap. Water was provided *ad libitum*. Mice were removed to their home cages after 4-8 hours on collection and urine was aspirated from plastic wrap and collected in microcentrifuge tubes. Urine was centrifuged at 150xg at room temperature for 5 minutes to remove cell debris and subsequently filtered using 0.22 um Ultrafree-MC devices (Millipore-Sigma) according to manufacturer’s instructions. Filtrate was further diafiltrated using Amicon Ultra 2mL 100K (Millipore-Sigma) according to manufacturer’s instructions (40 minutes concentration time at 4000xg). Filtrate was resuspended in an identical volume of DMEM as collected urine. 50% confluent NMuMG cells in a 24-well plate were infected for 1 h at 37°C with 150 μl / well of freshly diafiltrated urine from positive and negative mice harvested on days 18, 19, 50, 114, 127, and 169 d.p.i. After infection, complete media was added and cells were incubated at 37°C. Cells were passaged when confluent and the percentage of cytopathic effect (CPE) was noted daily in increments of 25%. CPE was scored as the fraction of cells appearing refractile, floating, or dead. The four C57BL/6N mice used as mock-infected controls always tested negative both via PCR and CPE.

### Determining mutations in muPyV genomes found in female mice’s lung and tumor

DNA purified as described above was diluted 1:1000 in water before amplification. Full muPyV genomes from the female mouse FL’s lung and tumor were amplified by two PCR using the Phusion^®^ High-Fidelity DNA Polymerase (New England Biolabs), as recommended by the manufacturer. The primers used in the inverted PCR are NGS_Rev 5’-GAATATAGCTGAATACACAGTTTATTC-3’ and FullSeq4 5’-GTGAAATCCAACACCATGTG-3’ and the primers used in the second PCR are NGS_Fwd 5’-CATGGCCTCCCTCATAAGTT-3’ and FGAR 5’-CACAAACAGTATGGGCCCG-3’. Full muPyV genomes from the female mouse FR’s lung were amplified by inverted PCR using the Phusion^®^ High-Fidelity DNA Polymerase (New England Biolabs), as recommended by the manufacturer, and the primers FullBC2XhoI 5’- CTCGAGCGTGACCAGTTTGCTAGTGAG-3’ and FullBC2P 5’-ACCTCCTTCACAAGACCCTG-3’. The PCR products were cloned into the plasmid pUC19 at SmaI and Sanger sequenced.

### Illumina NextSeq library preparation

#### Enrichment PCR

The Illumina NextSeq SR75 library was performed essentially as previously described for urine and virus stock samples (36). For DNA samples from urine, typically 20ul of eluted DNA solution was used per PCR reaction. For DNA samples from whole blood and virus stock, a maximum of 4.77 x 10^5^ copies of muPyV genomes (estimated by qPCR) was used per PCR reaction. For DNA samples purified from organs, we also used a maximum of 4.77 x 10^5^ copies of muPyV genomes but within a limit of 3.1μg of total DNA per reaction (i.e., the maximum quantity of DNA with no inhibitory effect on the PCR). The number of enrichment PCR cycles varied based on the copies of muPyV genomes used in the reaction.

To amplify DNA from organs and whole blood, 50μl of the PCR reaction contained ≤4.77 x 10^5^ copies of muPyV genome and ≤3.1μg of total DNA per PCR reaction, 0.3μM of each primer, 1X of KAPA HiFi HotStart Ready Mix (Kapa Biosystems). After an initial denaturation step for 3 min at 95°C, the amplification was performed by 22-35 cycles of 20 sec at 98°C (ramp 2°C/sec), 15 sec at 56°C (ramp 2°C/sec), 15 sec at 72°C (ramp 2°C/sec) followed by a final extension step for 2 min at 72°C. PCR reactions were stored immediately at -20°C until use. The presence of a specific amplification product was checked on a 2% agarose gel.

### Indexing PCR, amplicons pooling and quality controls

The indexing PCR was performed as previously described (36). In brief, the enrichment PCR products were used in two separate indexing PCR reactions per sample to amplify a 75 bp fragment encompassing the barcode region of the muPyVs genome using staggered indexing primers. The two indexing PCR reactions were merged, gel purified, quantified, normalized and pooled together to constitute the final library, as described previously (36). The final library was sent to the Genomic Sequencing and Analysis Facility of the University of Texas at Austin for a Bioanalyzer (Agilent) quality control prior to the Illumina NextSeq SR75 sequencing run. To increase diversity, PhiX DNA was also included (∼5% to the first run and ∼34% to the second run). The accession number for these data deposited at SRA is: PRJNA791340.

### Extracting barcode sequences

This was performed as previously described (36). In brief, barcodes were extracted from FASTQ files off the sequencer using Cutadapt (80), with the default error allowance of 10%, using linked adapters (requiring both adapters to flank the barcode). Relative abundance was determined by pulling the raw counts and processing them using R with the tidyverse packages (81).

### Clustering and quantifying barcodes

Based on our previous work (36), barcodes were clustered using Starcode (82) for message-passing clustering, using a maximum Levenshtein distance of 3 and the default cluster ratio of 5, with a cutoff applied to include only the 99% most abundant reads. This resulted in an estimate of 4012 unique barcodes in our initial input barcoded stock virus. To determine abundance of barcodes in both urine and tissues, we associated every barcode in a sample to the nearest stock barcode based on Levenshtein distance. If there was no stock barcode within a distance of 3, we dropped that sample barcode (distance 3 was a cutoff). If a sample barcode was close to n different stock barcodes, we assigned 1/n of the count to each of the nearest barcodes. For example, if a sample barcode was closest to 2 different stock barcodes, we assigned 1/2 of the count to one stock barcode, and 1/2 to the other. Then the counts were normalized to take into account that some samples had more muPyV bulk DNA (viral genomes) than others. For each sample and each barcode, we calculated what fraction that barcode count was of the total and then multiplied the fraction by the total number of estimated genome copies in the biological sample

### Randomized barcode *in silico* experiment

In order to validate the observed patterns of unique barcodes and account for potential background noise, we generated an *in silico* control by randomly shuffling the nucleotides of each sample barcode. These shuffled sequences were mapped to stock barcodes using the same threshold of Levenshtein distance ≤ 3 using R. For each sample, we counted the number of unique barcodes detected after mapping to the stock library for both the original and shuffled barcodes. The *in silico* control provided a baseline for distinguishing meaningful barcode patterns from random noise. This experiment was repeated using 5 different shufflings and the background baseline remained consistent across the trials.

### Analysis of barcode overlap between urine and tissues

To assess whether the most abundant shed viruses originated from the tissue reservoirs, we analyzed the overlap between the top 5% of barcodes in urine and tissues for each mouse. The overlap between these barcode sets was assessed using a contingency table that categorized barcodes into four groups: (1) present in the top 5% of both urine and tissue, (2) present only in the top 5% of urine, (3) present only in the top 5% of tissue, and (4) present in neither category. Fisher’s exact test was performed to evaluate the statistical significance of the overlap.

### Plotting data

Plots for number of unique barcodes, enrichment over background, similarity of temporally adjacent time points, changes in diversity over time (donut plots and entropy), shedding patterns via ridge plots, rank plots, GC content, length distribution of barcodes, Fisher’s exact test and correlation analyses (83) were generated using R with the tidyverse packages (81). We also used the ggridges and corrplot packages in R for generating the ridge plots and correlation plots respectively. The gridExtra and patchwork packages in R were used to combine plots for panel view. For correlation analyses, the Spearman correlation coefficient of the relative abundance of barcodes was calculated for each tissue of a given mouse using R. *Note, for determining the length of the barcode insert, we acknowledge that a caveat of this analysis is that it cannot identify virus genomes that may have entirely lost the barcode insert because these would not have been detected with the RT primer used in this analysis, which is designed to prime to a portion of the exogenous barcode insert.

Code available at: https://github.com/ChrisSullivanLab/Shedding-dynamics-of-a-DNA-virus

### Cosine similarity

Cosine similarity is a measure between -1 and 1 of the similarity between two sequences of numbers, where the proportions, rather than the absolute values of the numbers, are compared. For our purposes, the number of sequences mapping to a particular barcode represents the levels of the barcodes in a sample. More precisely, cosine similarity is the cosine of the angle between two vectors in an *n*-dimensional space, where for our purposes *n* is the number of distinct barcodes, and the coordinates of the vector for a sample are the levels of each barcode. If the levels of barcodes of one sample are simply a multiple of the levels for another sample, the cosine similarity between the two is 1. As applied to the barcode data, where the levels of barcodes are always non-negative, the cosine similarity must be between 0 and 1. We plotted a measure of cosine similarity between temporally adjacent time points as a measure of how the relatedness of the relative amounts of shed barcode patterns change over time.

### Diversity of viral genomes in samples

Donut charts are used to show the percentage representation of barcodes in each sample. Alternating shades of gray distinguish distinct barcodes, and the top 10 most abundant individual barcodes (“top 10” determined by the greatest amount of a barcode shed by an individual mouse over all timepoint tested) are indicated in a unique color.

To quantify the diversity of barcode repertoires over time, we utilized diversity indexes. We used the "true diversity", or Hill number *^q^D*, with *q* = 1 chosen so as not to prefer either abundant or rare barcodes. For *q* = 1, this is the exponential of the empirical Shannon entropy, also known as the Shannon-Wiener index (84). Additionally, we also counted how many barcodes it takes in each sample to account for 75% of the total barcode expression. Both measures of diversity are plotted as the percent change relative to the barcode repertoire shed at day 1.

## Author contributions

Designed wet bench experiments: SB, CSS

Performed wet bench experiments: SB

Analyzed data: SB, BMG, AM, CSS

Performed computational experiments: AM

Formatted data figure compilation: SB, BMG, AM, CSS

Generated Illumina libraries: SB

Developed and applied computational approaches: BMG, AM

Interpreted data: SB, BMG, AM, CSS

Managed project and procured funding: CSS

All authors assisted in the writing and editing of the manuscript

## Supplemental Figure Legends

**Figure S1. Barcoded polyomaviruses replicate infectious virus similar to wildtype virus.** Virus replication curves were plotted. A. NMuMG cells infected with WT muPyV (red) or barcoded muPyV (blue) at M.O.I. of 5. B. RPTEC infected with WT BKPyV (blue) or either of two different barcoded BKPyV libraries (red or green) at M.O.I. of 1. Cell-associated virus (muPyV) and supernatant (BKPyV) were harvested at multiple time post-infection (p.i.), in duplicate, and virus concentration was determined by immunofluorescence microscopy for VP1 viral proteins. This analysis shows similar kinetics of infectious virus production and confirms no obvious defect in virus replication caused by the barcode inserts.

**Figure S2. Ridge plots of the 10 most abundant barcodes.** A. Shown in color is the abundance of each of the top 10most shed barcodes in urine for each mouse (“top 10” determined by the greatest amount of a barcode shed at any single time point). The gray shaded area represents total bulk shed viral DNA at a particular time point post-infection. The height of an individual peak on the vertical axis correlates to the relative linear abundance of each barcode. The horizontal axis corresponds to different timepoints post-infection. B. Shows the sum total of the top 10 individual abundant shed barcodes for each mouse in gold (Note: this panel is identical to Figure 7).

**Figure S3. Mean GC content of shed barcodes does not change substantially during the course of infection.** A. GC content for the bulk of all barcodes **s**hed at each time point. B. GC content for the top 10 most shed barcodes for each mouse (“top 10” determined by the greatest amount of a barcode shed at any single time point). Neither panel shows overt trends towards altered nucleotide composition.

**Figure S4. Length of top 10 most abundant shed barcodes for each mouse.** The length of the barcodes for the top 10 most abundant shed barcodes (“top 10” determined by the greatest amount of a barcode shed at any single time point). Most barcodes retain an insert length of 12 nucleotides, as expected from the library design.

**Figure S5. Total amounts of muPyV DNA in organs.** Plotted are muPyV genomes equivalent determined by qPCR and normalized per µg of total DNA purified from organs or per µl of whole blood.

**Figure S6.** Statistical significance of barcode overlap between the top 5% of barcodes in urine and tissue samples across animals. The y-axis represents the −log10 transformed p-values from Fisher’s exact test, indicating the strength of association between barcode presence in urine and tissue. Asterisks denote significance levels: *p<0.05, **p<0.01, ***p<0.001, ****p<0.0001, and *****p<0.00001.

**Figure S7. GC Content of top 10 most abundant barcodes in tissues in each animal.** Shown is the GC content of the top 10 most abundant barcodes with abundance determined as the sum total of genome equivalents for each barcode in all tissues assayed for that animal (“top 10” determined by the greatest amount of a barcode shed at any single time point).

**Figure S8. Abundant barcodes in kidney are more abundantly shed in urine.** Shown are the top 10 most abundant barcodes detected in each organ of a given mouse and their rank in urine that was collected on the final day before sacrifice (“top 10” determined by the greatest amount of a barcode in any tissue for an individual mouse). In 3 of the 4 mice, abundant barcodes in the kidney are clearly also more abundant in urine. No other organ or tissue displayed such a strong signature consistent with shed viruses deriving from the kidney.

## Supplemental Tables

**Table S1: Virus stocks titer (IU/mL) as determined by immunofluorescence assay.**

**Table S2: Urine from infected mice contains infectious muPyV as shown by 100% CPE on NMuMG cells**

**Table S3- Median rank of top 10 most abundant in urine or tissue for each mouse.**

**Table S4: Mutations in muPyV genomes found in female mice’s lung and tumor. Table S5: Staggered indexing primers.**

## References

1. Amato KA, Haddock LA 3rd, Braun KM, Meliopoulos V, Livingston B, Honce R, Schaack GA, Boehm E, Higgins CA, Barry GL, Koelle K, Schultz-Cherry S, Friedrich TC, Mehle A. 2022. Influenza A virus undergoes compartmentalized replication in vivo dominated by stochastic bottlenecks. Nat Commun 13:3416.

2. Aliota MT, Dudley DM, Newman CM, Weger-Lucarelli J, Stewart LM, Koenig MR, Breitbach ME, Weiler AM, Semler MR, Barry GL, Zarbock KR, Haj AK, Moriarty RV, Mohns MS, Mohr EL, Venturi V, Schultz-Darken N, Peterson E, Newton W, Schotzko ML, Simmons HA, Mejia A, Hayes JM, Capuano S 3rd, Davenport MP, Friedrich TC, Ebel GD, O’Connor SL, O’Connor DH. 2018. Molecularly barcoded Zika virus libraries to probe in vivo evolutionary dynamics. PLoS Pathog 14:e1006964.

3. McCune BT, Lanahan MR, tenOever BR, Pfeiffer JK. 2020. Rapid Dissemination and Monopolization of Viral Populations in Mice Revealed Using a Panel of Barcoded Viruses. J Virol 94.

4. Varble A, Albrecht RA, Backes S, Crumiller M, Bouvier NM, Sachs D, García-Sastre A, tenOever BR. 2014. Influenza A virus transmission bottlenecks are defined by infection route and recipient host. Cell Host Microbe 16:691–700.

5. Weger-Lucarelli J, Garcia SM, Rückert C, Byas A, O’Connor SL, Aliota MT, Friedrich TC, O’Connor DH, Ebel GD. 2018. Using barcoded Zika virus to assess virus population structure in vitro and in Aedes aegypti mosquitoes. Virology 521:138–148.

6. Erickson AK, Jesudhasan PR, Mayer MJ, Narbad A, Winter SE, Pfeiffer JK. 2018. Bacteria Facilitate Enteric Virus Co-infection of Mammalian Cells and Promote Genetic Recombination. Cell Host Microbe 23:77–88.e5.

7. Pletnev AG, Maximova OA, Liu G, Kenney H, Nagata BM, Zagorodnyaya T, Moore I, Chumakov K, Tsetsarkin KA. 2021. Epididymal epithelium propels early sexual transmission of Zika virus in the absence of interferon signaling. Nat Commun 12:2469.

8. Riemersma KK, Jaeger AS, Crooks CM, Braun KM, Weger-Lucarelli J, Ebel GD, Friedrich TC, Aliota MT. 2021. Rapid evolution of enhanced Zika virus virulence during direct vertebrate transmission chains. J Virol 95:JVI.02218-20.

9. Sexton Nicole R., Bellis Eric D., Murrieta Reyes A., Spangler Mark Cole, Cline Parker J., Weger-Lucarelli James, Ebel Gregory D. 2021. Genome Number and Size Polymorphism in Zika Virus Infectious Units. J Virol 95:10.1128/jvi.00787-20.

10. Khanal S, Fennessey CM, O’Brien SP, Thorpe A, Reid C, Immonen TT, Smith R, Bess JWJ, Swanstrom AE, Del Prete GQ, Davenport MP, Okoye AA, Picker LJ, Lifson JD, Keele BF. 2019. In Vivo Validation of the Viral Barcoding of Simian Immunodeficiency Virus SIVmac239 and the Development of New Barcoded SIV and Subtype B and C Simian-Human Immunodeficiency Viruses. J Virol 94.

11. Fennessey CM, Pinkevych M, Immonen TT, Reynaldi A, Venturi V, Nadella P, Reid C, Newman L, Lipkey L, Oswald K, Bosche WJ, Trivett MT, Ohlen C, Ott DE, Estes JD, Del Prete GQ, Lifson JD, Davenport MP, Keele BF. 2017. Genetically-barcoded SIV facilitates enumeration of rebound variants and estimation of reactivation rates in nonhuman primates following interruption of suppressive antiretroviral therapy. PLoS Pathog 13:e1006359.

12. Marsden MD, Zhang T, Du Y, Dimapasoc M, Soliman MSA, Wu X, Kim JT, Shimizu A, Schrier A, Wender PA, Sun R, Zack JA. 2020. Tracking HIV Rebound following Latency Reversal Using Barcoded HIV. Cell Rep Med 1:100162.

13. Kim JT, Zhang T-H, Carmona C, Lee B, Seet CS, Kostelny M, Shah N, Chen H, Farrell K, Soliman MSA, Dimapasoc M, Sinani M, Blanco KYR, Bojorquez D, Jiang H, Shi Y, Du Y, Komarova NL, Wodarz D, Wender PA, Marsden MD, Sun R, Zack JA. 2022. Latency reversal plus natural killer cells diminish HIV reservoir in vivo. Nat Commun 13:121.

14. Krump NA, Liu W, You J. 2018. Mechanisms of persistence by small DNA tumor viruses. Curr Opin Virol 32:71–79.

15. Whitley RJ, Kimberlin DW, Roizman B. 1998. Herpes Simplex Viruses. Clin Infect Dis 26:541–553.

16. Wherry EJ, Ahmed R. 2004. Memory CD8 T-Cell Differentiation during Viral Infection. J Virol 78:5535–5545.

17. Speck SH, Ganem D. 2010. Viral latency and its regulation: lessons from the gamma-herpesviruses. Cell Host Microbe 8:100–15.

18. Benjamin TL. 2001. Polyoma Virus: Old Findings and New Challenges. Virology 289:167–173.

19. Howley PM, Livingston DM. 2009. Small DNA tumor viruses: Large contributors to biomedical sciences. Virology 384:256–259.

20. Ramqvist T, Dalianis T. 2010. Lessons from immune responses and vaccines against murine polyomavirus infection and polyomavirus-induced tumours potentially useful for studies on human polyomaviruses. Anticancer Res 30:279–84.

21. Swanson PA, Lukacher AE, Szomolanyi-Tsuda E. 2009. Immunity to polyomavirus infection: The polyomavirus–mouse model. Semin Cancer Biol 19:244–251.

22. Ramqvist T, Dalianis T. 2009. Murine polyomavirus tumour specific transplantation antigens and viral persistence in relation to the immune response, and tumour development. Semin Cancer Biol 19:236–243.

23. McCance DJ, Mims CA. 1979. Reactivation of polyoma virus in kidneys of persistently infected mice during pregnancy. Infect Immun 25:998–1002.

24. Moser JM, Lukacher AE. 2001. Immunity to Polyoma Virus Infection and Tumorigenesis. Viral Immunol 14:199–216.

25. Imperiale MJ, Jiang M. 2016. Polyomavirus Persistence. Annu Rev Virol 3:517–532.

26. Gardner SD, Field AM, Coleman DV, Hulme B. 1971. New human papovavirus (B.K.) isolated from urine after renal transplantation. Lancet Lond Engl 1:1253–7.

27. Padgett BL, Walker DL, ZuRhein GM, Eckroade RJ, Dessel BH. 1971. Cultivation of papova-like virus from human brain with progressive multifocal leucoencephalopathy. Lancet Lond Engl 1:1257–60.

28. Haycox CL, Kim S, Fleckman P, Smith LT, Piepkorn M, Sundberg JP, Howell DN, Miller SE. 1999. Trichodysplasia Spinulosa – A Newly Described Folliculocentric Viral Infection in an Immunocompromised Host. J Investig Dermatol Symp Proc 4:268–271.

29. Gaynor AM, Nissen MD, Whiley DM, Mackay IM, Lambert SB, Wu G, Brennan DC, Storch GA, Sloots TP, Wang D. 2007. Identification of a Novel Polyomavirus from Patients with Acute Respiratory Tract Infections. PLoS Pathog 3:e64.

30. Feng H, Shuda M, Chang Y, Moore PS. 2008. Clonal Integration of a Polyomavirus in Human Merkel Cell Carcinoma. Science 319:1096–1100.

31. van der Meijden E, Janssens RWA, Lauber C, Bouwes Bavinck JN, Gorbalenya AE, Feltkamp MCW. 2010. Discovery of a New Human Polyomavirus Associated with Trichodysplasia Spinulosa in an Immunocompromized Patient. PLoS Pathog 6:e1001024.

32. DeCaprio JA, Garcea RL. 2013. A cornucopia of human polyomaviruses. Nat Rev Microbiol 11:264–76.

33. Mishra N, Pereira M, Rhodes RH, An P, Pipas JM, Jain K, Kapoor A, Briese T, Faust PL, Lipkin WI. 2014. Identification of a Novel Polyomavirus in a Pancreatic Transplant Recipient With Retinal Blindness and Vasculitic Myopathy. J Infect Dis 210:1595–1599.

34. Burke JM, Bass CR, Kincaid RP, Ulug ET, Sullivan CS. 2018. The Murine Polyomavirus MicroRNA Locus Is Required To Promote Viruria during the Acute Phase of Infection. J Virol 92.

35. Rowe WP. 1961. The epidemiology of mouse polyoma virus infection. Bacteriol Rev 25:18–31.

36. Blois S, Goetz BM, Bull JJ, Sullivan CS. 2022. Interpreting and de-noising genetically engineered barcodes in a DNA virus. PLOS Comput Biol 18:e1010131.

37. Dubensky TW, Villarreal LP. 1984. The primary site of replication alters the eventual site of persistent infection by polyomavirus in mice. J Virol 50:541–6.

38. Atkin SJL, Griffin BE, Dilworth SM. 2009. Polyoma virus and simian virus 40 as cancer models: history and perspectives. Semin Cancer Biol 19:211–217.

39. Fluck MM, Schaffhausen BS. 2009. Lessons in Signaling and Tumorigenesis from Polyomavirus Middle T Antigen. Microbiol Mol Biol Rev 73:542–563.

40. Zhou AY, Ichaso N, Adamarek A, Zila V, Forstova J, Dibb NJ, Dilworth SM. 2011. Polyomavirus middle T-antigen is a transmembrane protein that binds signaling proteins in discrete subcellular membrane sites. J Virol 85:3046–54.

41. Brostoff T, Dela Cruz FN, Church ME, Woolard KD, Pesavento PA. 2014. The raccoon polyomavirus genome and tumor antigen transcription are stable and abundant in neuroglial tumors. J Virol 88:12816–24.

42. Berger JR, Danaher RJ, Dobbins J, Do D, Miller CS. 2020. Dynamic expression of JC virus in urine and its relationship to serostatus. Mult Scler Relat Disord 41:101972.

43. Li R-M, Branton MH, Tanawattanacharoen S, Falk RA, Jennette JC, Kopp JB. 2002. Molecular identification of SV40 infection in human subjects and possible association with kidney disease. J Am Soc Nephrol JASN 13:2320–2330.

44. Murphy E, Vanicek J, Robins H, Shenk T, Levine AJ. 2008. Suppression of immediate-early viral gene expression by herpesvirus-coded microRNAs: Implications for latency. Proc Natl Acad Sci 105:5453–5458.

45. Speck SH, Ganem D. 2010. Viral latency and its regulation: lessons from the gamma-herpesviruses. Cell Host Microbe 8:100–15.

46. Knipe DM, Cliffe A. 2008. Chromatin control of herpes simplex virus lytic and latent infection. Nat Rev Microbiol 6:211–221.

47. Mellerick DM, Fraser NW. 1987. Physical state of the latent herpes simplex virus genome in a mouse model system: evidence suggesting an episomal state. Virology 158:265–75.

48. Stevens JG, Wagner EK, Devi-Rao GB, Cook ML, Feldman LT. 1987. RNA complementary to a herpesvirus alpha gene mRNA is prominent in latently infected neurons. Science 235:1056–9.

49. Umbach JL, Kramer MF, Jurak I, Karnowski HW, Coen DM, Cullen BR. 2008. MicroRNAs expressed by herpes simplex virus 1 during latent infection regulate viral mRNAs. Nature 454:780–783.

50. Garber DA, Schaffer PA, Knipe DM. 1997. A LAT-associated function reduces productive-cycle gene expression during acute infection of murine sensory neurons with herpes simplex virus type 1. J Virol 71:5885–93.

51. Mador N, Goldenberg D, Cohen O, Panet A, Steiner I. 1998. Herpes simplex virus type 1 latency-associated transcripts suppress viral replication and reduce immediate-early gene mRNA levels in a neuronal cell line. J Virol 72:5067–75.

52. Cliffe AR, Coen DM, Knipe DM. 2013. Kinetics of facultative heterochromatin and polycomb group protein association with the herpes simplex viral genome during establishment of latent infection. mBio 4:e00590–12.

53. Cliffe AR, Garber DA, Knipe DM. 2009. Transcription of the herpes simplex virus latency-associated transcript promotes the formation of facultative heterochromatin on lytic promoters. J Virol 83:8182–90.

54. Wang Q-Y, Zhou C, Johnson KE, Colgrove RC, Coen DM, Knipe DM. 2005. Herpesviral latency-associated transcript gene promotes assembly of heterochromatin on viral lytic-gene promoters in latent infection. Proc Natl Acad Sci U S A 102:16055–9.

55. Thompson RL, Sawtell NM. 2001. Herpes Simplex Virus Type 1 Latency-Associated Transcript Gene Promotes Neuronal Survival. J Virol 75:6660–6675.

56. Kwiatkowski DL, Thompson HW, Bloom DC. 2009. The polycomb group protein Bmi1 binds to the herpes simplex virus 1 latent genome and maintains repressive histone marks during latency. J Virol 83:8173–81.

57. Flores O, Nakayama S, Whisnant AW, Javanbakht H, Cullen BR, Bloom DC. 2013. Mutational inactivation of herpes simplex virus 1 microRNAs identifies viral mRNA targets and reveals phenotypic effects in culture. J Virol 87:6589–603.

58. Cai W, Astor TL, Liptak LM, Cho C, Coen DM, Schaffer PA. 1993. The herpes simplex virus type 1 regulatory protein ICP0 enhances virus replication during acute infection and reactivation from latency. J Virol 67:7501–12.

59. Pan D, Flores O, Umbach JL, Pesola JM, Bentley P, Rosato PC, Leib DA, Cullen BR, Coen DM. 2014. A Neuron-Specific Host MicroRNA Targets Herpes Simplex Virus-1 ICP0 Expression and Promotes Latency. Cell Host Microbe 15:446–456.

60. Arthur RR, Shah KV. 1989. Occurrence and significance of papovaviruses BK and JC in the urine. Prog Med Virol Fortschritte Med Virusforsch Progres En Virol Medicale 36:42–61.

61. Egli A, Infanti L, Dumoulin A, Buser A, Samaridis J, Stebler C, Gosert R, Hirsch HH. 2009. Prevalence of polyomavirus BK and JC infection and replication in 400 healthy blood donors. J Infect Dis 199:837–46.

62. Heritage J, Chesters PM, McCance DJ. 1981. The persistence of papovavirus BK DNA sequences in normal human renal tissue. J Med Virol 8:143–50.

63. Imperiale MJ, Major EO. 2007. Polyomaviruses, p. 2263–2298. *In* Knipe, DM, Howley, PM (eds.), Fields Virology, Fifth Edition. Lippincott Williams & Wilkins, Philadelphia.

64. Kwun HJ, Chang Y, Moore PS. 2017. Protein-mediated viral latency is a novel mechanism for Merkel cell polyomavirus persistence. Proc Natl Acad Sci U S A 114:E4040–E4047.

65. Cuddy SR, Cliffe AR. 2023. The Intersection of Innate Immune Pathways with the Latent Herpes Simplex Virus Genome. J Virol 97:e0135222.

66. Van der Sluis RM, Zerbato JM, Rhodes JW, Pascoe RD, Solomon A, Kumar NA, Dantanarayana AI, Tennakoon S, Dufloo J, McMahon J, Chang JJ, Evans VA, Hertzog PJ, Jakobsen MR, Harman AN, Lewin SR, Cameron PU. 2020. Diverse effects of interferon alpha on the establishment and reversal of HIV latency. PLoS Pathog 16:e1008151.

67. Ryabchenko B, Soldatova I, Šroller V, Forstová J, Huérfano S. 2021. Immune sensing of mouse polyomavirus DNA by p204 and cGAS DNA sensors. FEBS J 288:5964–5985.

68. Qin Q, Shwetank, Frost EL, Maru S, Lukacher AE. 2016. Type I Interferons Regulate the Magnitude and Functionality of Mouse Polyomavirus-Specific CD8 T Cells in a Virus Strain-Dependent Manner. J Virol 90:5187–5199.

69. Wu H-H, Li Y-J, Weng C-H, Hsu H-H, Chang M-Y, Yang H-Y, Yang C-W, Tian Y-C. 2023. Interferon-alpha and MxA inhibit BK polyomavirus replication by interaction with polyomavirus large T antigen. Biomed J 100682.

70. Assetta B, De Cecco M, O’Hara B, Atwood WJ. 2016. JC Polyomavirus Infection of Primary Human Renal Epithelial Cells Is Controlled by a Type I IFN-Induced Response. mBio 7.

71. An P, Sáenz Robles MT, Duray AM, Cantalupo PG, Pipas JM. 2019. Human polyomavirus BKV infection of endothelial cells results in interferon pathway induction and persistence. PLoS Pathog 15:e1007505.

72. Butic AB, Spencer SA, Shaheen SK, Lukacher AE. 2023. Polyomavirus Wakes Up and Chooses Neurovirulence. Viruses 15:2112.

73. van Aalderen MC, Heutinck KM, Huisman C, ten Berge IJM. 2012. BK virus infection in transplant recipients: clinical manifestations, treatment options and the immune response. Neth J Med 70:172–183.

74. Ambalathingal GR, Francis RS, Smyth MJ, Smith C, Khanna R. 2017. BK Polyomavirus: Clinical Aspects, Immune Regulation, and Emerging Therapies. Clin Microbiol Rev 30:503–528.

75. Polo C, Pérez JL, Mielnichuck A, Fedele CG, Niubo J, Tenorio A. 2004. Prevalence and patterns of polyomavirus urinary excretion in immunocompetent adults and children. Clin Microbiol Infect 10:640–644.

76. Hirsch HH, Steiger J. 2003. Polyomavirus BK. Lancet Infect Dis 3:611–623.

77. Hirsch HH, Randhawa PS, AST Infectious Diseases Community of Practice. 2019. BK polyomavirus in solid organ transplantation— Guidelines from the American Society of Transplantation Infectious Diseases Community of Practice. Clin Transplant 33:e13528.

78. Qin Q, Lauver M, Maru S, Lin E, Lukacher AE. 2017. Reducing persistent polyomavirus infection increases functionality of virus- specific memory CD8 T cells. Virology 502:198–205.

79. Peden KW, Pipas JM, Pearson-White S, Nathans D. 1980. Isolation of mutants of an animal virus in bacteria. Science 209:1392–6.

80. Martin M. 2011. Cutadapt removes adapter sequences from high-throughput sequencing reads. EMBnet.journal 17:10–12.

81. Wickham H, Averick M, Bryan J, Chang W, McGowan L, François R, Grolemund G, Hayes A, Henry L, Hester J, Kuhn M, Pedersen T, Miller E, Bache S, Müller K, Ooms J, Robinson D, Seidel D, Spinu V, Yutani H. 2019. Welcome to the Tidyverse. J Open Source Softw 4:1686.

82. Zorita E, Cuscó P, Filion GJ. 2015. Starcode: sequence clustering based on all-pairs search. Bioinforma Oxf Engl 31:1913–9.

83. Wei T, Simko V. 2021. R package “corrplot”: Visualization of a Correlation Matrix. https://github.com/taiyun/corrplot.

84. Tuomisto H. 2010. A consistent terminology for quantifying species diversity? Yes, it does exist. Oecologia 164:853–860.

